# Precursor peptide-targeted mining of more than one hundred thousand genomes expands the lanthipeptide natural product family

**DOI:** 10.1101/2020.03.13.990614

**Authors:** Mark C. Walker, Douglas A. Mitchell, Wilfred A. van der Donk

## Abstract

**Background:** Lanthipeptides belong to the ribosomally synthesized and post-translationally modified peptide group of natural products and have a variety of biological activities ranging from antibiotics to antinociceptives. These peptides are cyclized through thioether crosslinks and can bear other secondary post-translational modifications. While lanthipeptide biosynthetic gene clusters can be identified by the presence of characteristic enzymes involved in the post-translational modification of these peptides, locating the precursor peptides encoded within these clusters is challenging due to their short length and high sequence variability, which limits the high-throughput exploration of lanthipeptide precursor peptides. To address this challenge, we enhanced the predictive capabilities of Rapid ORF Description & Evaluation Online (RODEO) to identify all known classes of lanthipeptides.

**Results:** Using RODEO, we mined over 100,000 bacterial and archaeal genomes in the RefSeq database. We identified nearly 8,500 lanthipeptide precursor peptides. These precursor peptides were identified in a broad range of bacterial phyla as well as the Euryarchaeota phylum of archaea. Bacteroidetes were found to encode a large number of these biosynthetic gene clusters, despite making up a relatively small portion of the genomes in this dataset. While a number of these precursor peptides are similar to those of previously characterized lanthipeptides, even more were not, including potential antibiotics. Additionally, examination of the biosynthetic gene clusters revealed enzymes that install secondary post-translational modifications are more widespread than initially thought.

**Conclusion:** Lanthipeptide biosynthetic gene clusters are more widely distributed and the precursor peptides encoded within these clusters are more diverse than previously appreciated, demonstrating that the lanthipeptide sequence-function space remains largely underexplored.

---

Ribosomally synthesized and post-translationally modified peptides (RiPPs) are an expanding group of natural products [1]. Lanthipeptides are among the most studied RiPPs and have a diverse array of structures and biological activities, including antibiotic [2-5], anti-fungal [6], anti-HIV [7, 8], and antinociceptive [9, 10] activities. These peptidic natural products are characterized by the presence of macrocycles formed via thioether crosslinks between amino acid residue side chains, termed lanthionines or methyllanthionines [11]. Lanthipeptides are synthesized from a genetically encoded precursor peptide, generically named LanA, which can be divided into two portions; an N-terminal leader region, involved in recognition by the biosynthetic machinery, and a C-terminal core region, which is post-translationally modified. The essential enzymes in lanthipeptide biosynthesis dehydrate select serine and threonine residues in the core region to form dehydroalanine and dehydrobutyrine residues, respectively, and then catalyze the conjugate addition of cysteine thiols onto the resulting double bonds to form the lanthionine or methyllanthionine crosslinks (Fig. 1A). Lanthipeptides can be divided into four classes based on the essential biosynthetic enzymes [11]. In class I lanthipeptides, separate proteins carry out the dehydration (LanB) [12] and cyclization (LanC) [13] reactions. LanB enzymes activate serine and threonine residues by glutamylation in a tRNA-dependent manner and produce the dehydrated residues through beta-elimination of glutamate. In classes II-IV, a single protein carries out dehydration and cyclization (LanM, LanKC, and LanL, respectively) (Fig. 1B) [11]. The C-terminal cyclization domains of LanMs, LanKCs, and LanLs are homologous to LanC cyclases; however, the LanKC cyclization domain lacks the zinc-binding residues that are conserved in the other cyclases [14]. The LanM dehydratase domain is related to lipid kinases [15] whereas the LanKC and LanL proteins catalyze dehydration using dedicated kinase [16] and lyase domains. The latter are related to OspF, a phosphothreonine lyase from certain pathogenic proteobacteria [17]. Beyond dehydratases and cyclases, lanthipeptide biosynthetic gene clusters (BGCs) often encode transporters and proteases to remove the leader peptide (LanT/LanP) and sometimes additional enzymes that further decorate lanthipeptides with secondary modifications [11, 18].

**Fig. 1.**
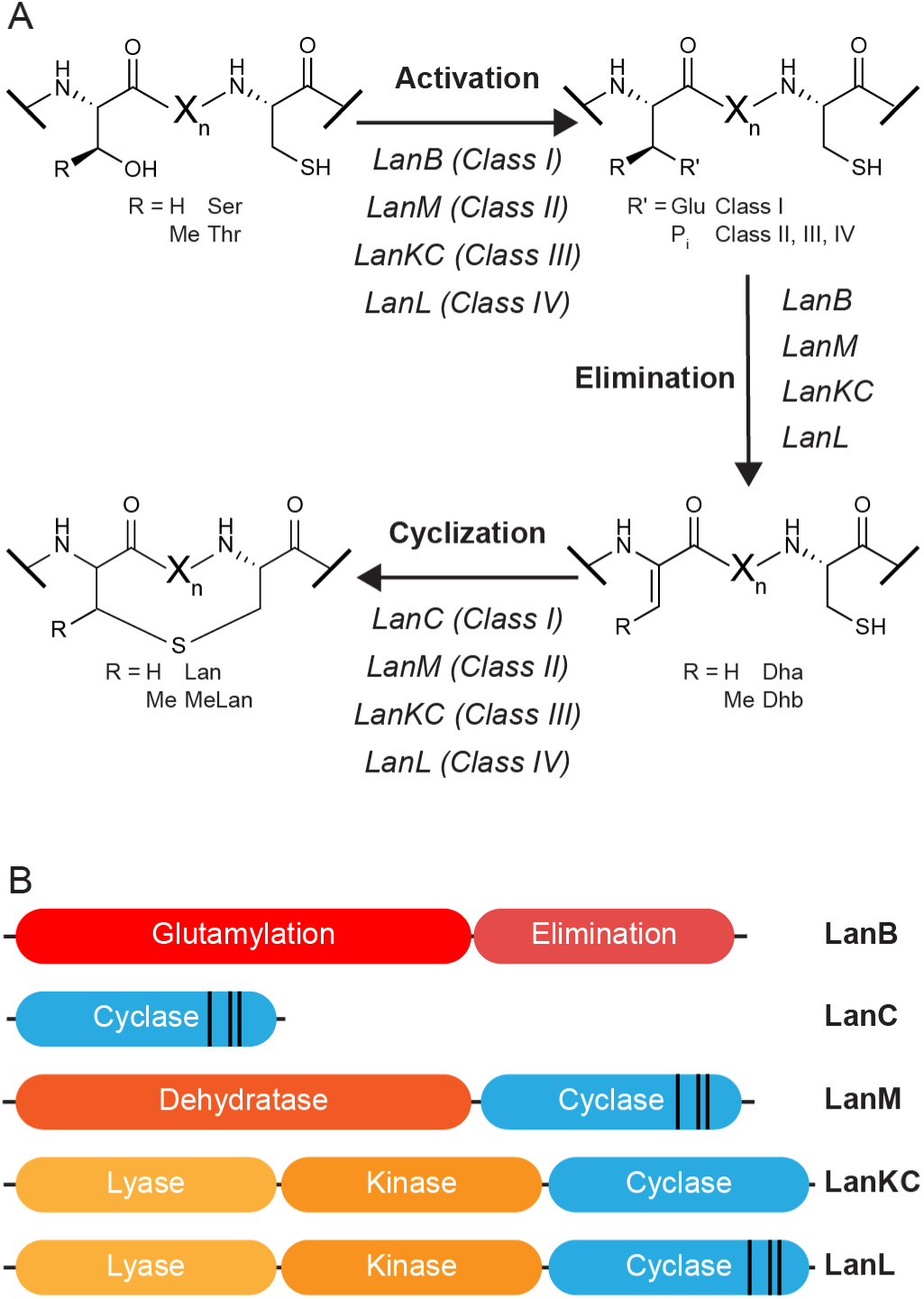
Biosynthesis of lanthipeptides. A) Installation of lanthionine or methyllanthionine thioether crosslinks in the four different classes of lanthipeptides. Dha: dehydroalanine, Dhb: dehydrobutyrine, Lan: lanthionine, MeLan: methyllanthionine. B) Domain structure of the enzymes that install the thioether crosslinks in the different classes of lanthipeptides. The black lines in the cyclase domains represent the location of the zinc-binding residues.

Genome-mining studies based on these enzymes have revealed that lanthipeptide BGCs are distributed widely across bacterial phyla [19-29]. Despite the success in bioinformatically identifying likely lanthipeptide BGCs, it has been an outstanding challenge to perform high-throughput analysis of the precursor peptides encoded in these gene clusters. Because of the short length of the genes encoding LanAs, they are often not annotated as genes and their variability renders identifying new precursors through homology searching challenging. To address this problem, we have expanded Rapid ORF Description & Evaluation Online (RODEO) [30] to predict lanthipeptide precursor peptides and mined the bacterial and archaeal genomes in the RefSeq database for new lanthipeptide natural products.

## Results

### Identification of potential lanthipeptide biosynthetic gene clusters

Potential lanthipeptide BGCs were identified by searching the non-redundant RefSeq database (release 93) with the LANC_like (PF05147) hidden Markov model (HMM) from the Protein family (Pfam) database [31], as this domain is shared among currently known classes of lanthipeptides (Figure 1B). This search resulted in 12,572 proteins with LanC-like domains. The genomic context of these proteins was then examined to assign the clusters to the four separate lanthipeptide classes. If any of the proteins encoded in the seven genes upstream or downstream from the LanC-domain containing protein matched the Pfam HMMs for a LanB (PF04738 and PF14028), the cluster was categorized as class I. If the encoded proteins matched the Pfam HMM for the dehydratase domain of a LanM (PF13575), the cluster was categorized as class II. If the protein containing the LanC-like domain also matched the Pfam HMM for a protein kinase (PF00069), the cluster was categorized as class III or class IV. Classes III and IV were then separated using custom HMMs to distinguish LanKCs (class III) from LanLs (class IV). If none of the encoded proteins matched with these Pfam HMMs, the cluster was categorized as unclassified. This sorting resulted in 2,753 putative class I lanthipeptide BGCs, 3,675 class II BGCs, 2,377 class III BGCs, 815 class IV BGCs, and 2,919 unclassified sequences. With the exception of 33 putative class II BGCs from Archaea, lanthipeptide BGCs were exclusively identified in bacteria. Of the unclassified proteins, 1,146 are likely not within lanthipeptide BGCs as these proteins are more similar to other proteins, such as endogluconases [13]. In another 381 cases of unclassified proteins, the gene encoding the protein is within 3 kb of the beginning or end of a sequencing contig, suggesting incomplete data on the BGC. Intriguingly, a number of the remaining 1,392 unclassified proteins are located within BGCs that encode proteins often associated with RiPP biosynthesis, such as ABC transporters and proteases, suggesting these clusters are potentially involved in the biosynthesis of an as-of-yet uncharacterized class of RiPP.

### Identification of precursor peptides

Having identified potential lanthipeptide BGCs, we set out to identify the cognate precursor peptide(s). The DNA sequence spanning the cluster was searched for potential open reading frames (ORFs) beginning with an ATG, TTG, or GTG start codon. Potential ORFs that encoded peptides within the expected length range for LanAs (30 – 120 amino acids) and not located entirely within an annotated ORF were identified for scoring. A random subset consisting of 20% of the BGCs for each class were manually examined and the identified peptides were annotated as a precursor peptide or not. Next, 2,458 features were calculated for these peptides (Supplementary Figure S1, Additional File 1) and ANOVA was used to identify the features that were most significantly different (p-value < 0.05) between high-confidence precursor peptides and likely unrelated peptides. These features were then calculated for the entire set of potential precursor peptides for each class, and the manually annotated peptides were used as a training set for support vector machine (SVM) classification of the peptides as precursor or not. The SVM classification, the presence of sequence motifs in the leader peptide, and other features were used in the RODEO framework to identify potential precursor peptides (Supplementary Tables S1-4, Additional File 1).

This approach resulted in the identification of 8,406 precursor peptides (Additional File 2). Of these putative LanAs, 2,698 (32% of the total) were from class I BGCs, 3,002 (36%) were from class II BGCs, 2,304 (27%) were from class III BGCs, and 401 (5%) were from class IV BGCs. Approximately 24% of class I precursors, 17% of class II precursors, 55% of class III precursors, and 86% of class IV precursors were not annotated as genes. The most abundant, ungapped sequence motifs from the leader and core regions of each class were identified using Multiple Em for Motif Elicitation (MEME) (Supplementary Figure S2, Additional File 1) [32]. None of the leader peptide motifs were shared among lanthipeptide classes, which was expected given the differences in the respective lanthionine synthases. Interestingly, the most abundant core motifs from each class were also restricted to that class. For example, the nisin/gallidermin lipid II-binding motif SxxxTP(G/S)C [33] is only found in class I precursors and the mersacidin lipid II-binding motif TxTxEC [34, 35] is only found in class II precursors. Examining these sequence motifs also reveals that in addition to the long-recognized FxLD sequence motif in the leader peptides of class I LanAs [36], a number of class I LanAs from Bacteroidetes have a LxLxKx_5_L motif instead. Many of the leader peptides that contain this motif end with a Gly-Gly sequence, and a C39-family Cys protease that removes leader peptides at GG sites [37, 38] is often encoded in the corresponding clusters. This GG leader motif has previously only been observed in class II [39] and III LanAs [40]. With the identification of these class I LanAs, approximately one third of all LanAs have a GG motif separating the leader and core peptide. Double-Gly motif leader peptides are also a common occurrence in other RiPP classes [1, 41]. Other frequently observed leader peptide sequence motifs are the (E/D−8)(L/M−7) motif in class II [42]and the LxLQ motif in class III lanthipeptide precursors (Supplementary Figure S2) [43]. Additional less frequent motifs that have not been experimentally investigated are depicted in Supplementary Figure S2.

A sequence similarity network analysis [44] (Fig. 2) reveals that the precursor peptides tend to cluster into families by lanthipeptide class and by taxonomic phylum (Fig. 2 and Supplementary Figure S3, Additional File 1). Even though a number of these families include lanthipeptides that have been characterized, as indicated by the representative lanthipeptides shown (Supplementary Table S5, Additional File 1), most families lack a characterized member, highlighting the scope of lanthipeptide sequence space that remains to be studied. In this work, we have labeled the precursor families by a Roman numeral indicating lanthipeptide class and an increasing Arabic number from left to right and top to bottom in the order generated by the Organic Layout of Cytoscape [45]. Several of the uncharacterized families, including I 8, I 13, II 18, and II 32, appear to contain lipid II-binding motifs (Supplementary Figures S4-7, Additional File 1) and are likely antibiotics. The four largest class I families (I 1-4) are from Actinobacteria and do not have a characterized member. Their core peptides contain a highly conserved Asp residue that is of particular note because the corresponding BGCs contain an *O*-methyltransferase (PF01135) and the conserved Asp is likely post-translationally modified [46]. A number of the class II families, such as II 2, II 13, II 17, and II 26 have conserved leader peptides and non-conserved core peptides. The leader peptides from families II 2, II 13, II 17, II 25, and II 29 belong to the nitrile hydratase leader peptide family of leader peptides, whereas the leader peptides from family II 26 belong to the Nif11 family of leader peptides [41]. The precursor peptides in family II 26 are from Cyanobacteria, however the prochlorosin lanthipeptides are not among them [47]. The prochlorosin precursor peptides are located in a smaller cluster, which does not represent the actual size of this family of precursors as many of them are encoded in genes located distantly from their cognate LanM in the genome [47, 48]. We suggest the name cyanotins for this family of RiPPs that are made from highly diverse core peptides, some of which lack Cys and hence cannot be precursors to lanthipeptides.

**Fig. 2.**
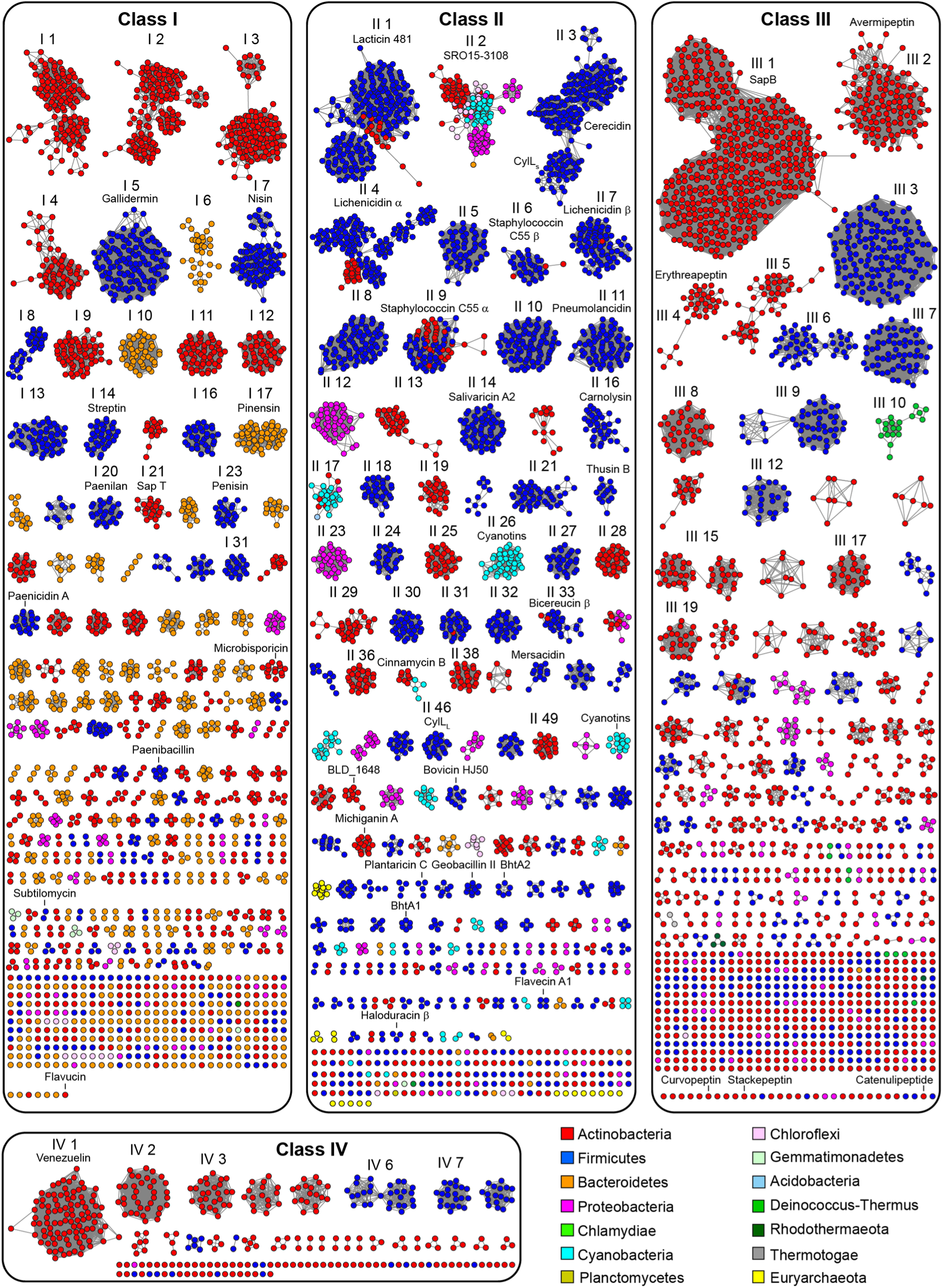
Sequence similarity networks [44] of precursor peptides. Clusters of precursor peptides with 20 or more members are numbered and sequence logos for these clusters are presented in Supplementary Figure S3. Clusters with characterized members as determined by using BAGEL4 [27] and the MIBiG repository [85] (Supplementary Table S5) are labeled by a selected member.

### Other enzymes in lanthipeptide biosynthetic gene clusters

Other proteins present in the BGCs were characterized by searching the Pfam database of HMMs. Examining the most abundant proteins that hit at least one Pfam family reveals lanthionine synthases, proteases, ABC transporters, and transcriptional regulators (Supplementary Tables S6-9, Additional File 1). A number of class I BGCs contain split LanB enzymes that contain the glutamylation and elimination domains on separate polypeptides, as is seen in the biosynthesis of the lanthipeptide pinensin [6], as well as the thiopeptide family of RiPPs [49]. Other class I BGCs contain a full length LanB and a protein homologous to the LanB elimination domain. These proteins are also homologous to the enzyme in thiopeptide biosynthesis that catalyzes a formal [4+2] cycloaddition to install a substituted pyridine or (dehydro)piperidine moiety [12, 50-52]. Accordingly, it is an intriguing possibility that these domains catalyze a post-translational modification other than elimination. These standalone elimination domain proteins are also often fused to protein-L-isoaspartate *O*-methyltransferase (PCMT or PIMT, PF01135) family proteins and, in turn, many BGCs have these *O*-methyltransferases as standalone proteins. Notably, these elimination domain proteins and methyltransferases are nearly exclusively limited to class I BGCs (Supplementary Table S10, Additional File 1).

Enzymes that are among the most abundant in one class of lanthipeptide BGCs are generally also present in the other classes, if at lower abundance (Supplementary Table S10 and Figure S8, Additional File 1). For example, flavoprotein family enzymes, which have been shown to catalyze oxidative decarboxylation of the C-terminus of some lanthipeptides (LanDs) [53-57], halogenation of amino acid side chains [55], and oxidation of the sulfur in lanthionine crosslinks [58], are among the most abundant enzymes in class I BGCs but are present in class II and III BGCs as well. Likewise, NAD(P)H-dependent FMN reductase family enzymes, such as those that catalyze the reduction of dehydro amino acid side chains to form D-amino acid residues (LanJ_B_s) [59, 60], are among the most common tailoring enzymes in class II BGCs and are present in class I and III BGCs. Another enzyme family, the zinc-dependent dehydrogenases, have been demonstrated to carry out the same overall reaction [61], and members of this family are present in all four classes of lanthipeptide BGCs (Supplementary Table S11). To date, the installation of D-amino acids has only been observed in class II lanthipeptides, but these reductases and dehydrogenases suggest these structures may also be present in class I, III, and IV lanthipeptides, or alternatively, these enzymes may catalyze a new post-translational modification. Some BGCs from all four classes of lanthipeptides encode a short chain dehydrogenase. This family of enzymes has been shown to install an N-terminal lactate moiety [62], although this modification has thus far only been observed in class I lanthipeptides. To date, no secondary post-translational modifications have been reported for class IV lanthipeptides; however, a number of these clusters contain genes encoding FAD-dependent oxidoreductases, glycosyltransferases, and acetyltransferases. Thus, tailoring may occur for the products of these clusters, or alternatively, the genes encoding these other enzymes may not be part of the gene clusters.

Many BGCs appear to encode enzymes that are less widely distributed but may carry out rare post-translational modifications (Supplementary Table S11 and Figure S9, Additional File 1). For example, some class I, II, and III lanthipeptide BGCs contain a YcaO family protein (PF02624), members of which catalyze modification to the amide backbone [63]. Moreover, a number of BGCs for all four classes of lanthipeptides encode polyketide or fatty acid biosynthetic machinery, as in the recently reported class III lipolanthine [56], or non-ribosomal peptide biosynthetic machinery. Enzymes from other families, such as radical SAM (PF04055), cytochrome P450 (PF00067), and α-ketoglutarate-dependent oxygenases (PF03171), are present in lanthipeptide BGCs and may catalyze the installation of additional secondary modifications.

### Phylogenetic distribution of lanthipeptide biosynthetic clusters

Lanthipeptide biosynthetic enzymes were identified in a wide range of bacterial phyla, with the majority (within currently sequenced genomes) in Actinobacteria (Fig. 3A). The distribution of these proteins across phyla is inconsistent for the different classes of lanthipeptides (Fig. 3B). Nearly a quarter of the class I LanCs were identified in Bacteroidetes, despite their genomes making up a relatively small portion of those in the data set (Supplementary Figure S10, Additional File 1). This distribution suggests further genome sequencing efforts of Bacteroidetes may uncover additional novel lanthipeptide BGCs. At present, only the pinensins have been isolated from this phylum [6]. LanMs were the only lanthipeptide biosynthetic enzymes identified in Cyanobacteria. The majority of LanKCs and LanLs are from Actinobacteria and Firmicutes (Figure 2); however, no members of these class III or class IV lanthipeptides from Firmicutes have been characterized.

**Fig. 3.**
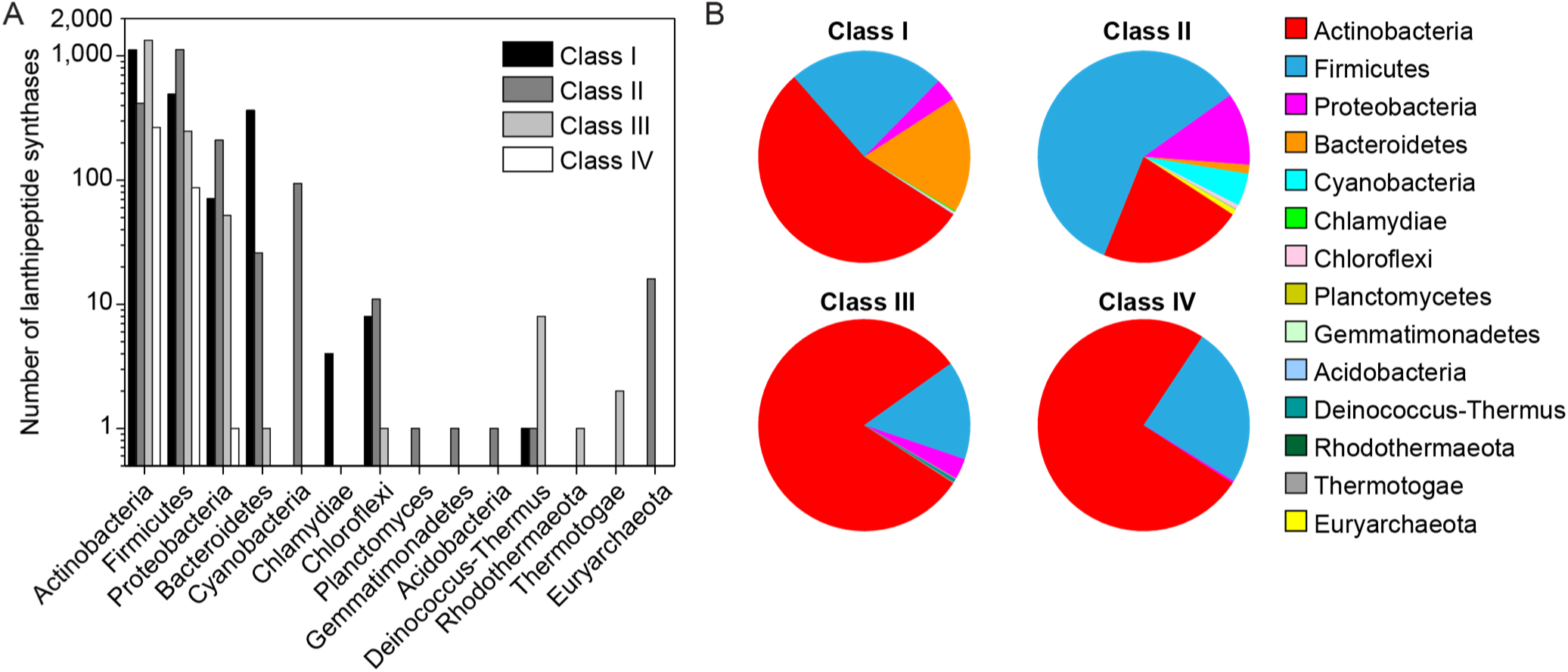
Phylogenetic distribution of lanthipeptide biosynthetic gene clusters. A) The distribution of lanthipeptide BGCs in bacterial and archaeal phyla. B) The distribution of bacterial and archaeal phyla among the four classes of lanthipeptide BGCs.

A phylogenetic tree of LanCs and LanC-like domains reveals clades corresponding to the class of lanthipeptide and then sub-clades of bacterial phyla (Fig. 4). This topology suggests the divergence of the lanthipeptide classes is ancient and supports the hypothesis that the different lanthipeptide classes through convergent evolution [64]. Inclusion of human LanC-like proteins on the tree shows that they fall into the class IV clade, which is made up of proteins with LanC-like domains linked to a kinase domain (Supplementary Figure S11, Additional File 1). Notably, human LanC-like proteins bind to kinases in various cell lines [65]. Some exceptions to grouping by class are observed, such as a class I LanCs from Bacteroidetes that appear to be related to the LanC-like domains of class II LanMs from Firmicutes. The precursors associated with these LanCs fall into cluster I 17, which includes the antifungal lanthipeptide pinensin [6]. Furthermore, a group of the LanC-like domains of LanMs from Actinobacteria are related to LanCs from the same phylum with the precursors associated with these LanMs falling in cluster II 28. By examining the sub-clades for each class, it appears that LanCs are more related to homologs from their own phylum than to other classes. Such relationships could suggest inter-phyla horizontal gene transfer occurs less frequently with class I BGCs. This lower incidence of horizontal gene transfer may be related to the use of, and selectivity for, glutamyl-tRNA by LanBs in class I lanthipeptide biosynthesis and the previously observed incompatibility of non-native tRNA^Glu^ sequences with activity by LanB enzymes [66]. To further explore the possibility of horizontal gene transfer, an analysis of the %GC content of the lanthipeptide BGCs versus the %GC content of the entire bacterial or archaeal genome was performed. Generally, these two values are in good agreement; however, the standard deviation of the differences between the %GC content of the genome and the cluster is smaller for class I lanthipeptide BGCs than for class II and III, potentially supporting a lower incidence of horizontal gene transfer of class I clusters (Supplementary Figure S12, Additional File 1).

**Fig. 4.**
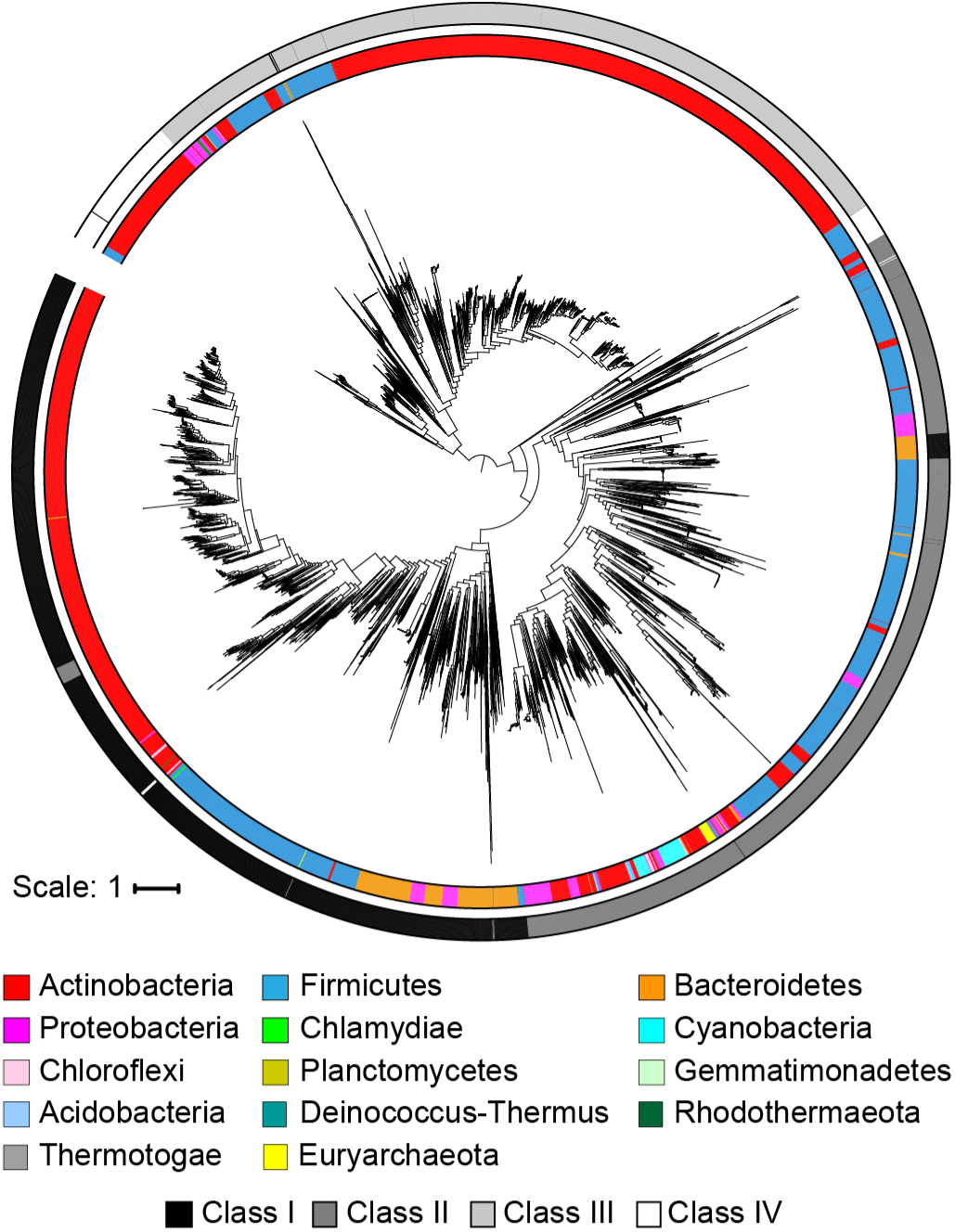
Phylogenetic analysis of LanC-like domains. A midpoint rooted approximately maximum likelihood phylogenetic tree of LanC-like domains from lanthipeptide biosynthetic enzymes. The inner ring is colored according to prokaryotic phylum and the outer ring is colored according to lanthipeptide class.

## Discussion

In this work, we improved the ability of RODEO to predict precursor peptides for all four classes of lanthipeptides. This expanded functionality facilitated the mining of more than one hundred thousand bacterial and archaeal genomes for the ability to produce lanthipeptides. These studies revealed that lanthipeptide BGCs are more broadly distributed than previously appreciated, with a large number of class I lanthipeptides in Bacteroidetes and the presence of class III and IV lanthipeptides in Firmicutes. Examining the precursor peptides encoded in the gene clusters revealed that the majority of lanthipeptide natural product families have not been characterized, including a number that are likely antibiotics because of lipid II-binding motifs. As delineated below, several new insights have been revealed through this bioinformatics study.

As in a previous study that focused on Actinobacteria [23], the most common lanthipeptide precursor family when analyzing all currently available genomes from different phyla (III 1) is the morphogenic SapB peptide involved in sporulation [67]. The third- and fourth-most abundant precursor families (II 3 and II 4) comprise single- and two-component lanthipeptides in which two structurally dissimilar lanthipeptides exert synergistic bioactivity, with the individual peptides usually having low or no activity [68]. The fourth most abundant family (II 4) includes the two-component lanthipeptide lichenicidin α (II 4) [28, 69, 70], which is primarily found in Firmicutes. However, a small number of family members is found in Actinobacteria, a phylum not previously reported to produce two-component lanthipeptides. Unexpectedly, the precursors of the partner lanthipeptide that would make up the two-component system (lichenicidin β, II 7) are less abundant and are almost exclusively encoded by Firmicutes, while the putative partners of the actinobacterial lichenicidin α–like precursors are in a different family (II 49; Fig. 5). Precursors related to another two-component lanthipeptide, staphylococcin C55, share a similar bifurcated distribution. The precursor of staphylococcin C55 α is a member of family II 9, which is comprised of peptides encoded in Firmicutes and Actinobacteria. However, the precursor of staphylococcin C55 β in Firmicutes is part of family II 14, whereas the putative partners of the Actinobacterial staphylococcin C55 α precursors are in family II 19. The α peptides of currently investigated two-component lanthipeptides are involved in lipid II-binding. The resulting complex is believed to serve as a binding site for the β peptides, which results in pore formation in the bacterial membranes [69, 71, 72]. The difference in ring topology between the β peptides of Firmicutes and Actinobacteria potentially suggests a different mode of action for the Actinobacterial β peptides, or that a different structure is required to form pores in the membranes of the target bacteria for lanthipeptide producers that live in different ecological niches.

**Fig. 5.**
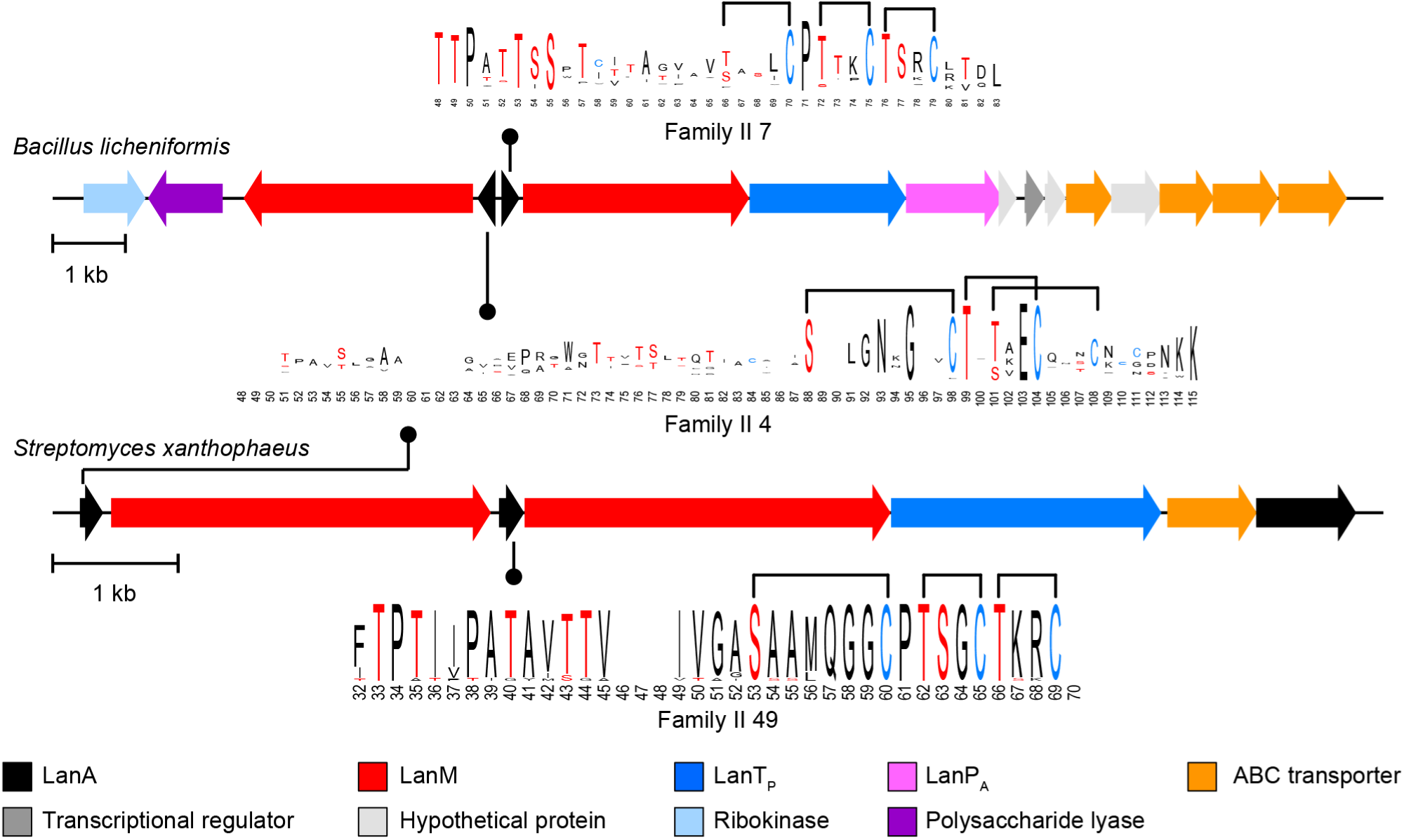
Representative diagrams for two-component lanthipeptide BGCs that share an α precursor peptide (from family II 4) with β peptides from families II 7 or II 49. Sequence logos for the predicted core peptides are shown as well. In the sequence logos, residues that can be dehydrated (i.e. Ser and Thr) are shown in red and Cys residues are shown in blue and predicted cyclization patterns are shown. The predicted cyclization pattern for family II 49 is based on similar positioning of modifiable residues in family II 7.

The precursor peptides of the third most abundant family of peptides (II 3) have sequence homology with the CylL_S_′′ peptide that together with CylL_L_′′ makes up the enterococcal cytolysin. This two-component lantibiotic lyses bacterial and mammalian cells [73], and epidemiological studies have shown a clear correlation between the presence of the cytolysin biosynthetic gene locus and hospital-acquired infections [74]. However, even more noticeable than for the lichenicidin system discussed above, the precursor peptides of the CylL_L_′′ peptides (II 30) are much less abundant than the precursor peptides for CylL_S_′′ (Fig. 2). Genes encoding these CylL_S_-like peptides are found mostly in species of *Bacillus* and sometimes in *Staphylococcus aureus*. Previous studies noted these peptides in *B. cereus* [21, 75] and several members have been isolated and shown to display antimicrobial activity without the need of a partner peptide.

The most abundant class I precursor peptides in Actinobacteria (I 1-4) are encoded in BGCs that contain the previously mentioned ortholog of a protein isoaspartate methyltransferase as well as a fully conserved Asp in the core peptide (Fig. 6A). This methyltransferase is the highest co-occurring protein with lanthionine biosynthetic enzymes (Additional File 1: Supplementary Table S9) even though it is only found in class I lanthipeptide BGCs. Recently, it was shown that one such enzyme methylated the conserved Asp in a precursor peptide of family I 2 encoded in *Streptomyces olivaceus* NRRL B-3009. Methylation led to the formation of a succinimide that was hydrolyzed to a mixture of aspartate and isoaspartate. The mature form of the natural product remains unknown [46].

**Fig. 6.**
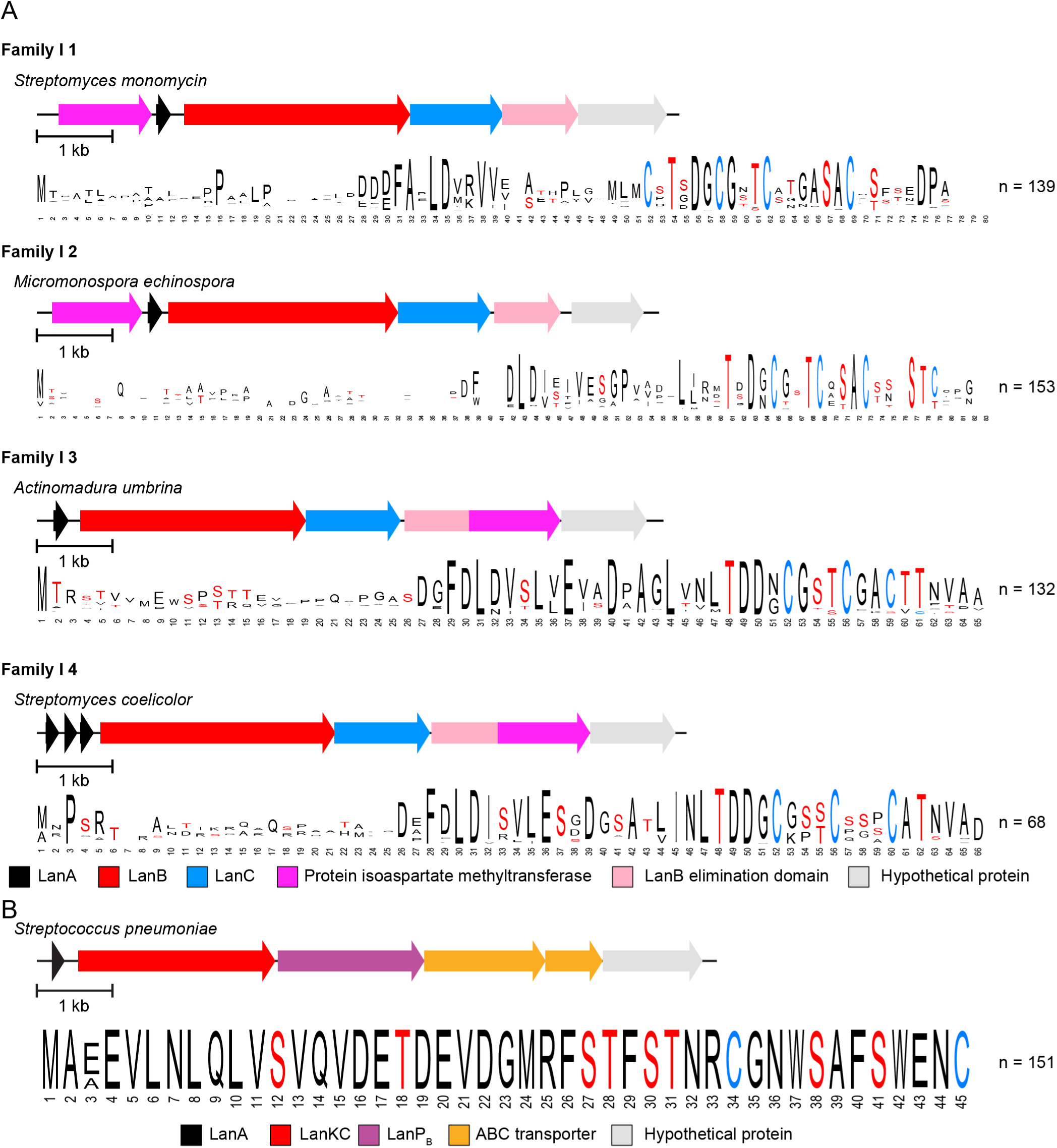
A. Representative diagrams for BGCs containing precursor peptides from families I 1, I 2, I 3, and I 5 along with sequence logos for those precursor families. B. A representative diagram for BGCs containing a precursor peptide from family III 3 along with the sequence logo for that family. LanP_B_ is a prolyl oligopeptidase family protease.

The sixth most abundant precursor family (III 3) consists of class III lanthipeptides encoded by Firmicutes; currently class III peptides have only been isolated from Actinobacteria. At present, the organisms encoding this lanthipeptide family are restricted to *Streptococcus pneumoniae*. The precursor peptides display identical core sequences (Fig. 6B) that contain the characteristic Sx_2_Sx_3_C motif that in some class III lanthipeptides give rise to a lanthionine between the first Ser and last Cys of the motif, and in other peptides yield labionin [10, 40, 67, 76] (Fig. 6B). Unlike class III lanthipeptide gene clusters from Actinobacteria that usually do not contain a protease [77], the III 3 clusters encode for a prolyl oligopeptidase (we propose the name LanP_B_ for these with the name LanP_A_ given to the previously characterized group of subtilisin-like leader peptide proteases [11]) as well as a CAAX family protease [78]. The smaller III 6 family is also found in Firmicutes but the phylogenetic distribution is more varied as these BGCs are found in different *Bacillus* and *Staphylococcus* species. Like class III BGCs from Actinobacteria, they do not encode a conserved protease, but they do encode an ABC-type transporter.

Generally speaking, a particular family of lanthipeptides is usually only produced by a single phylum, but some exceptions are notable. The lacticin 481 family is mostly produced by Firmicutes, but some Actinobacteria also encode members. Nisin, a commercially used food preservative, is almost exclusively encoded in Firmicutes genomes, but at present a single Actinobacterium (*Nocardia vaccinii* NBRC 15922) encodes this antimicrobial agent. After finding an example of nisin in an Actinobacteria we manually examined the genomic context of highly similar LanB and LanC enzymes. Nisin-like precursor peptides were identified in these BGCs, however, they were encoded more than seven genes away from the *lanC* gene and therefore fell outside of the genome neighborhood analyzed in this study. Some lanthipeptides seem to have distributed nearly equally between Firmicutes and Actinobacteria, such as the two-component lantibiotic lacticin 3147, as well as the II 2 family which is found roughly equally in three phyla, Firmicutes, Actinobacteria, and Cyanobacteria. At present, the molecular structures of members of this family have yet to be reported. Examination of the Pfam families of other enzymes encoded in the BGCs revealed that some of the tailoring enzymes that were previously thought to be limited to a single lanthipeptide class are in fact distributed among multiple, if not all, classes of lanthipeptides.

## Conclusions

The current comprehensive analysis of bacterial and archaeal genomes for the presence of *lanC*-like genes with the development of methodology to characterize the cognate precursor peptides reveal the diversity of the lanthipeptide family of RiPPs and the extent to which a large portion of chemical space remains to be explored. The current study will facilitate prioritization of genome-mining studies for novel structures, new synergistic lanthipeptide pairs, or lanthipeptides from genera currently not known to produce such compounds.

## Methods

### Bioinformatic mining for lanthipeptides

The non-redundant protein, GenBank, and nucleotide records for bacteria and archaea in the RefSeq collection were downloaded from NCBI in May of 2019. The non-redundant protein records were searched with the LanC-Like Pfam HMM (PF05147.12) [79] using HMMER3 [80] with the default settings. The GenBank and nucleotide records were parsed using the list of LanC-Like proteins. Proteins encoded within seven ORFs upstream and downstream of the LanC-Like protein were annotated by searching the Pfam HMM database using HMMER with an E-value cutoff of 1×10^−5^. Gene clusters that encoded proteins that matched with the Lant_dehydr_N Pfam HMM (PF04738.13) and the Lant_dehydr_C Pfam HMM (PF14028.6) were classified as class I clusters. Gene clusters that encoded a protein that matched with the DUF4135 Pfam HMM (PF13575.6) were classified as class II clusters. Gene clusters that encoded a protein that matched both the LANC_Like Pfam HMM and the Pkinase Pfam HMM (PF00069.25) were classified as class III or class IV clusters. Gene clusters that did not encode proteins that matched these HMMs were discarded as unclassified. Custom HMMs were developed for class III and class IV LanC-Like domains. The sequences of representative LanKCs and LanLs were aligned with Clustal-Omega [81], and these alignments were manually truncated to include only the LanC-like domain. HMMER was then used to generate HMMs from these alignments (Additional Files 3 and 4). The LanC-like proteins in the class III and IV gene clusters were then searched against these custom HMMs, and classified as class III if the E-value for the match with the class III HMM was lower than that for the class IV HMM, and as class IV if the E-value for the match with the class IV HMM was lower than that for the class III HMM. Additionally, the DNA sequence spanning the most upstream ORF and most downstream ORF was translated into all potential ORFs with ATG, GTG, or TTG start codons. Potential ORFs that were longer than 120 amino acids or shorter than 30 amino acids were discarded, as were any that were encoded entirely within an annotated gene. Additionally, any ORF not encoding a Cys was discarded as it could not be a potential lanthipeptide precursor. Finally, to reduce redundancy, the longest of the remaining ORFs with the same stop codon coordinates were retained for further analysis.

### Scoring of potential precursor peptides

Leader peptide motifs were identified in the ORFs identified above using the MEME bioinformatics application. The leader-core boundary was then estimated by searching the ORF following the leader peptide motif for GG, GA, or S/T(x)_2-7_C and setting the core region as starting immediately following a GG or GA motif or 1 residue before a S/T(x)_2-7_C as long as that motif was more than 10 residues from the end of the ORF. If multiple of these motifs were identified, the one allowing the longest core region was used as the boundary. If none of these motifs were present or were present within 10 residues of the C-terminus, the C-terminal half of the ORF was used as the core. If a leader peptide motif was not identified, the same analysis was performed from the beginning of the ORF. Finally, if no Cys was present in the estimated core region, the ORF was discarded as not a lanthipeptide precursor. Features were then calculated for the potential core peptide, SVM classification was performed, and the potential precursor peptides were scored according to the rubrics in Supplementary Tables S1 – S4.

### Phylogenetic analysis of LanC and LanC-like domains

LanC and LanC-like domain containing proteins from clusters encoding likely precursor peptides per the analysis above, were retrieved. LanM, LanKC, and LanL enzymes were aligned separately using Clustal-Omega, manually truncated to their LanC-like domains, and then unaligned. Then an alignment of LanCs and LanC-like domains was constructed using Clustal-Omega and manually edited to remove large gaps. This alignment was used to calculate an approximately maximum likelihood phylogenetic tree using FastTree [82]. The tree was then visualized using the Interactive Tree of Life [83].

### Precursor sequence logos

Likely precursor peptides were aligned using Clustal-Omega, and that alignment was used to generate sequence logos using WebLogo [84].

## List of Abbreviations

BGC: biosynthetic gene cluster,
HMM: hidden Markov model,
LanA: lanthipeptide precursor peptide;
LanB: class I lanthipeptide dehydratatse;
LanC: class I lanthipeptide cyclase
LanKC: class III lanthipeptide synthetase
LanL: class IV lanthipeptide synthetase
LanM: class II lanthipeptide synthetase;
MEME: Multiple Em for Motif Elicitation;
ORF: open reading frame.
Pfam: protein family;
RODEO: Rapid ORF Description & Evaluation Online;
RiPP: ribosomally synthesized and post-translationally modified peptide;
SVM: support vector machine.

## Declarations

### Ethics approval and consent to participate

Not applicable

### Consent for publication

Not applicable

### Availability of data and material

All biosynthetic gene clusters and precursor peptides identified in this study are available in Additional file

2. Genomes used in this study are available from NCBI in RefSeq release 93. The software used to perform the genome-mining studies is available at https://github.com/mcwalker-group/reimagined-octo-funicular.

### Author contributions

M.C.W., D.A.M., and W.A.V conceived of the study. M.C.W. performed the genome mining. All authors drafted the manuscript and have read and approved the final manuscript.

## Acknowledgements

We acknowledge technical assistance from Bryce Kille in implementing the lanthipeptide precursor peptide identification modules in RODEO.

## Additional Files

Additional File 1.

**Supplementary Figure S1**. Features calculated to score precursor peptides; **Supplementary Table S1**. Features and scoring for class I precursors; **Supplementary Table S2**. Features and scoring for class II precursors; **Supplementary Table S3**. Features and scoring for class III precursors; **Supplementary Table S4**. Features and scoring for class IV precursors; **Supplementary Figure S2**. Sequence motifs present in more than 100 lanthipeptide precursor peptides; **Supplementary Figure S3**. Sequence similarity network of predicted precursor peptides with permissive similarity cutoff; **Supplementary Table S5**. Location of top BLAST hits of known lanthipeptides from the MIBiG and BAGEL databases in the sequence similarity networks presented in Figure 2; **Supplementary Figure S4**. Sequence logos generated from alignments of class I precursor peptides in clusters with 20 or more members; **Supplementary Figure S5**. Sequence logos generated from alignments of class II precursor peptides in clusters with 20 or more members; **Supplementary Figure S6**. Sequence logos generated from alignments of class III precursor peptides in clusters with 20 or more members; **Supplementary Figure S7**. Sequence logos generated from alignments of class IV precursor peptides in clusters with 20 or more members; **Supplementary Table S6**. Twenty most abundant proteins in class I BGCs that belong to at least one Pfam; **Supplementary Table S7**. Twenty most abundant proteins in class II BGCs that belong to at least one Pfam; **Supplementary Table S8**. Twenty most abundant proteins in class III BGCs that belong to at least one Pfam; **Supplementary Table S9**. Twenty most abundant proteins in class IV BGCs that belong to at least one Pfam; **Supplementary Table S10**. Distribution of Pfams that are in the 20 most abundant protein families in one class among the other three classes; **Supplementary Figure S8**. Example biosynthetic gene clusters encoding the enzymes in Table S10; **Supplementary Table S11**. Distribution of select Pfam protein families from BGCs; **Supplementary Figure S9**. Example biosynthetic gene clusters encoding the enzymes in Table S11; **Supplementary Figure S10**. Phylogenetic distribution of genomes in the dataset; **Supplementary Figure S11**. A approximately maximum likelihood midpoint rooted phylogenetic tree of LanC and LanC-like domains including human LanC-like proteins; **Supplementary Figure S12**. GC content of clusters versus genomes.

Additional File 2.

Excel File containing precursor peptides identified in this study.

Additional File 3.

Custom HMM for LanKC LanC-like domains.

Additional File 4.

Custom HMM for LanL LanC-like domains.

