## Additional file 1 for "Precursor peptide-targeted mining of more than one hundred thousand genomes expands the lanthipeptide natural product family"

| | |
|---|---|
| Mark C. Walker | 0000-0003-3942-003X |
| Douglas A. Mitchell | 0000-0002-9564-0953 |
| Wilfred A. van der Donk | 0000-0002-5467-7071 |


### Supplementary Figures and Tables

**Supplementary Figure S1.** Features calculated for precursor peptides 2
**Supplementary Table S1.** Features and scoring for class I precursors 3
**Supplementary Table S2.** Features and scoring for class II precursors 3
**Supplementary Table S3.** Features and scoring for class III precursors 3
**Supplementary Table S4.** Features and scoring for class IV precursors 3
**Supplementary Figure S2.** Sequence motifs in precursor peptides 4
**Supplementary Figure S3.** Sequence similarity network of precursor peptides 5
**Supplementary Table S5.** Location of characterized lanthipeptides in sequence similarity network 6
**Supplementary Figure S4.** Sequence logos of class I precursor families 7
**Supplementary Figure S5.** Sequence logos of class II precursor families 9
**Supplementary Figure S6.** Sequence logos of class III precursor families 12
**Supplementary Figure S7.** Sequence logos of class IV precursor families 13
**Supplementary Table S6.** Abundant protein families in class I biosynthetic gene clusters 14
**Supplementary Table S7.** Abundant protein families in class II biosynthetic gene clusters 14
**Supplementary Table S8.** Abundant protein families in class III biosynthetic gene clusters 15
**Supplementary Table S9.** Abundant protein families in class IV biosynthetic gene clusters 15
**Supplementary Table S10.** Distribution of abundant protein families among lanthipeptide biosynthetic gene clusters 16
**Supplementary FigureS8.** Representative lanthipeptide biosynthetic gene clusters with abundant protein families 17
**Supplementary Table S11.** Distribution of select protein families in lanthipeptide biosynthetic gene clusters 25
**Supplementary Figure S9.** Representative lanthipeptide biosynthetic gene clusters with select protein families 26
**Supplementary Figure S10.** Phylogenetic distribution of genomes used in this study 32
**Supplementary Figure S11.** Phylogenetic tree of LanC-like proteins including human LanC-like proteins 33
**Supplementary Figure S12.** GC content of biosynthetic gene clusters and their cognate genomes 34
**References**. 35

**Supplementary Figure S1.** Features calculated to score precursor peptides. The DNA sequence encoding 7 genes upstream and downstream of a LanC-like domain containing protein was extracted and all potential open reading frames (ORFs) were identified. In the case that multiple potential ORFs shared a stop codon, the longest ORF within the expected range of LanA lengths was used. These ORFs were discarded if they were too long, too short, occurred entirely within an annotated gene, or did not contain a Cys residue. The remaining ORFs were then analyzed using FIMO [1] to identify conserved leader motifs. If the ORF contained a leader motif and has a GG, GA, or S/T $(x)_{2-7}$C motif downstream of the leader, but not within 10 residues of the end of the ORF, the approximate core was identified as starting immediately after the GG or GA motif or 1 reside before the S/T $(x)_{2-7}$C motif. If the ORF contained multiple GG or GA motifs, only the first one was considered, likewise with the S/T $(x)_{2-7}$C motif. If the ORF contained both a GG or GA motif and a S/T $(x)_{2-7}$C motif, the longer potential core was used. If the ORF did not contain a GG, GA, or S/T $(x)_{2-7}$C motif, or those motifs were within 10 residues of the end of the ORF, the C-terminal half of the peptide was identified as the potential core. If no leader peptide motif was identified in the ORF, the same analysis was performed starting from the beginning of the ORF instead of after the leader motif. Finally, if the predicted core did not contain a Cys residue, the ORF was discarded. The given features were then calculated for each potential ORF.

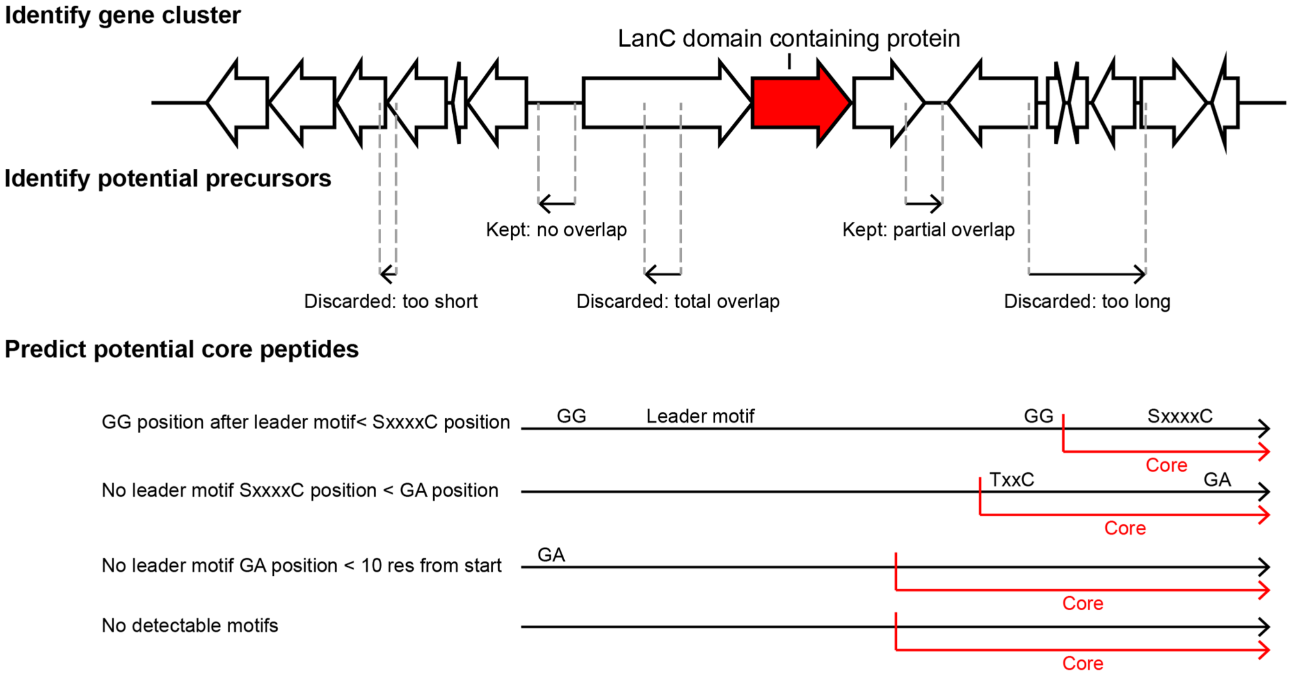


**Score potential core peptides**

Molecular weight of core
Isoelectric point of core
Number of each amino acid in core
Number of Ser + Thr in core
Number of Ser + Thr + Cys in core
Number of Charged residues in core
Number of positive charges in core
Number of negative charges in core
Net charge of core
Number of polar residues in core
Number of aliphatic residues in core
Number of aromatic residues in core

Core fraction that is Cys
Core fraction that is Ser + Thr
Core fraction that is Ser + Thr + Cys
Core fraction that is charged residues
Core fraction that is positive residues
Core fraction that is negative residues
Core fraction that is polar residues
Core fraction that is aliphatic residues
First fifth of precursor fraction S+T
First fifth of precursor fraction S+T+C
First fifth of precursor fraction C
Second fifth of precursor fraction S+T

Second fifth of precursor fraction S+T+C
Second fifth of precursor fraction C
Third fifth of precursor fraction S+T
Third fifth of precursor fraction S+T+C
Third fifth of precursor fraction C
Fourth fifth of precursor fraction S+T
Fourth fifth of precursor fraction S+T+C
Fourth fifth of precursor fraction C
Fifth fifth of precursor fraction S+T
Fifth fifth of precursor fraction S+T+C
Fifth fifth of precursor fraction C

Number of amino acid pairs

```
AA, AC, AD, ..., AW, AY   AxA, AxC, AxD, ..., AxW, AxY ... AxxxxxxA, AxxxxxxC, AxxxxxxD, ..., AxxxxxxW, AxxxxxxY
CA, CC, CD, ..., CW, CY   CxA, CxC, CxD, ..., CxW, CxY ... CxxxxxxA, CxxxxxxC, CxxxxxxD, ..., CxxxxxxW, CxxxxxxY
:                         :                                :
:                         :                                :
WA, WC, WD, ..., WW, WY   WxA, WxC, WxD, ..., WxW, WxY ... WxxxxxxA, WxxxxxxC, WxxxxxxD, ..., WxxxxxxW, WxxxxxxY
YA, YC, YD, ..., YW, YY   YxA, YxC, YxD, ..., YxW, YxY ... YxxxxxxA, YxxxxxxC, YxxxxxxD, ..., YxxxxxxW, YxxxxxxY
```

**Supplementary Table S1.** Features and scoring for class I precursors. ORFs were identified as precursors if their score was over 10. SVM, support vector machine.

| Feature | Score |
|---|---|
| SVM classification | 5 |
| Presence of Class I leader peptide MEME motif | 5 |
| Core pI less than 9 | 2 |
| 2 or more Cys in core | 2 |
| Leader has KLxLxK MEME motif and ends in GG sequence | 1 |

**Supplementary Table S2.** Features and scoring for class II precursors. ORFs were identified as precursors if their score was over 10.

| Feature | score |
|---|---|
| SVM classification | 5 |
| Presence of Class II leader peptide MEME motif | 5 |
| 2 or more Cys in core | 2 |
| Hits a Pfam* hidden Markov model | 5 |
| Precursor ends with sequence KRC | 4 |

*PF08130.1, PF04604.12, PF14867.5, PF16934.4, PF02979.15, PF07862.10

**Supplementary Table S3.** Features and scoring for class III precursors. ORFs were identified as precursors if their score was over 10.

| Feature | score |
|---|---|
| SVM classification | 5 |
| Presence of Class III leader peptide MEME motif | 5 |
| 2 or more Cys in core | 2 |
| Has SxxSxxxC motif | 1 |
| Has SxxSxxC motif | 1 |
| Core pI is between 3 and 9 | 1 |

**Supplementary Table S4.** Features and scoring for class IV precursors. ORFs were identified as precursors if their score was over 10.

| Feature | score |
|---|---|
| SVM classification | 5 |
| Presence of Class IV leader peptide MEME motif | 5 |
| 2 or more Cys in core | 2 |
| Precursor ends with sequence GCD | 1 |
| Precursor ends with sequence LGS | 1 |

**Supplementary Figure S2.** Sequence motifs present in more than 100 lanthipeptide precursor peptides. Some of the conserved leader peptide motifs have been shown to be important for the cognate lanthipeptide synthases (FxLD in class I [2]; E/D(-8)L/M(-9) in class II [3]; LxLQ in class III [4], and $Lx_2LPE$ in class IV [5]). Lipid II-binding motifs, which arise for likely antibiotic lanthipeptides are underlined in red.

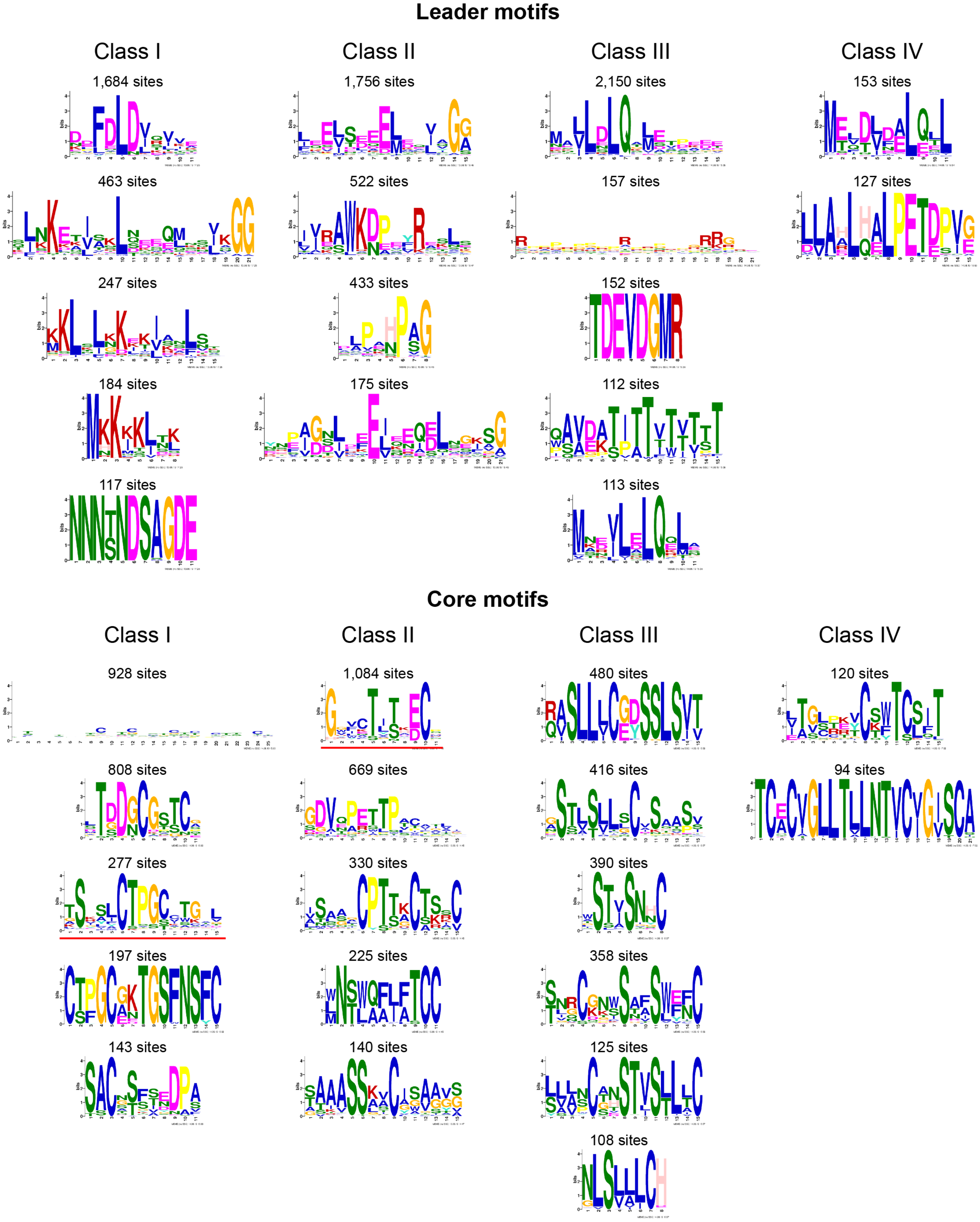

**Supplementary Figure S3.** Sequence similarity network of predicted precursor peptides generated with the Enzyme Function Initiative-Enzyme Similarity Tool [6] with permissive similarity cutoff (alignment score: 6; equivalent to an expectation value of $10^{-6}$) and visualized in Cytoscape [7].

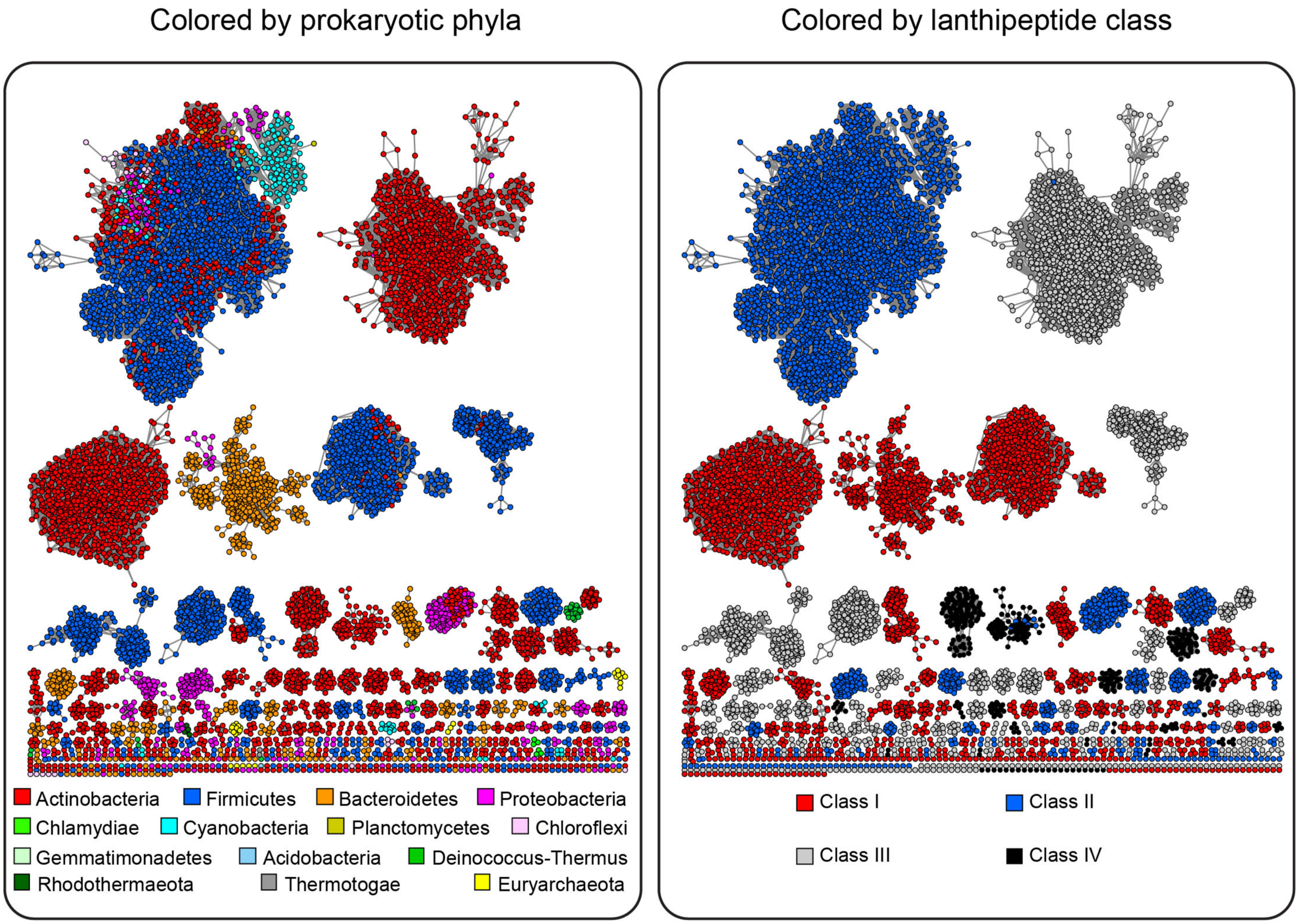

**Supplementary Table S5.** Location of top BLAST hits of known lanthipeptides from the MIBiG [8] and BAGEL [9] databases in the sequence similarity networks presented in Figure 2. If the precursor is located in a cluster with more than 20 members, the identifier of that cluster is given. If the precursor is located in a cluster with 20 or fewer members, the label from Figure 2 is given. If the precursor occurred only once, it is annotated as a singleton.

| Known Lanthipeptide | Cluster |
|---|---|
| AmfS | III 1 |
| BacCH91 | I 5 |
| BhtA1 | BhtA1 |
| BhtA2 | BhtA2 |
| Bicereucin_beta | II 33 |
| BLD_1648 | BLD_1648 |
| Bovicin_HJ50 | Bovicin HJ50 |
| BsaA2 | I 5 |
| Carnolysin_A1 | II 16 |
| Carnolysin_A2 | II 16 |
| Catenulipeptide | singleton |
| Cerecidin | II 3 |
| Cinnamycin_B | Cinnamycin B |
| ClylL | II 46 |
| ClylS | II 3 |
| Curvopeptin | singleton |
| Entianin | I 7 |
| Epidermin | I 5 |
| Ericin_A | I 7 |
| Ericin_S | I 7 |
| Erythreapeptin | III4 |
| Flavecin_A1 | Flavecin A1 |
| Flavucin | Flavucin |
| Gallidermin | I 5 |
| Geobacillin_I | I 7 |
| Geobacillin_II | Geobacillin II |
| Griseopeptin | III1 |
| Haloduracin_alpha | II 4 |
| Haloduracin_beta | Haloduracin_beta |
| Informatipeptin | III2 |
| Lichenicidin_VK21_A1 | II 4 |
| Lichenicidin_VK21_A2 | II 7 |
| Macedocin | II 1 |
| Macedovicin | Bovicin HJ50 |
| McdA1 | II 1 |
| Mersacidin | Mersacidin |
| Michiganin_A | Michiganin A |
| Microbisporicin | Microbisporicin |
| Mutacin_II | II 1 |
| Mutacin_K8 | II 1 |
| Nisin_A | I 7 |
| Nisin_O | I 7 |

| Known Lanthipeptide | Cluster |
|---|---|
| Nisin_U | I 7 |
| Nisin_Z | I 7 |
| Nukacin_ISK-1 | II 1 |
| Paenibacillin | Paenibacillin |
| Paenicidin_A | Paenicidin A |
| Paenicidin_B | Paenicidin A |
| Paenilan | I 20 |
| Penisin | I 23 |
| Pinensin | I 17 |
| Plantaricin C | PlantaricinC |
| Pneumolancidin_PldA2 | II 11 |
| Pneumolancidin_PldA3 | II 11 |
| Pneumolancidin_PldA4 | II 11 |
| Prochlorosin_1.1 | Prochlorosin 1.1 |
| Pseudomycoicidin | Geobacillin II |
| Ruminococcin_A | II 1 |
| SAL-2242 | III1 |
| Salivaricin_9 | II 1 |
| Salivaricin_A | II 14 |
| Salivaricin_G32 | II 1 |
| Sap_T | I 21 |
| SapB | III1 |
| Smb_A | BhtA2 |
| Smb_B | BhtA1 |
| SRO15-2212 | III1 |
| SRO15-3108 | II 2 |
| Stackepeptin_A | singleton |
| Stackepeptin_B | singleton |
| Stackepeptin_C | singleton |
| Stackepeptin_D | singleton |
| Staphylococcin_C55_alpha | II 9 |
| Staphylococcin_C55_beta | II 6 |
| Streptin | I 14 |
| Streptococcin_A-FF22 | II 1 |
| StreptococcinA-M49 | II 1 |
| Subtilin | I 7 |
| Subtilomycin | Subtilomycin |
| Suicin_3908 | Bovicin HJ50 |
| Suicin_65 | II 1 |
| Suicin_90-1330 | I 7 |
| Thermophilin_1277 | Bovicin HJ50 |
| Thusin_A | II 4 |
| Thusin_B | Thusin B |

**Supplementary Figure S4.** Sequence logos generated from alignments of class I precursor peptides in clusters with 20 or more members (in Figure 2) using WebLogo [10]. Conserved leader motifs that are shared among multiple clusters (from Figure S2) are boxed in gray and potential lipid II-binding motifs are overlined in red. Families with no previously characterized members are highlighted in yellow.

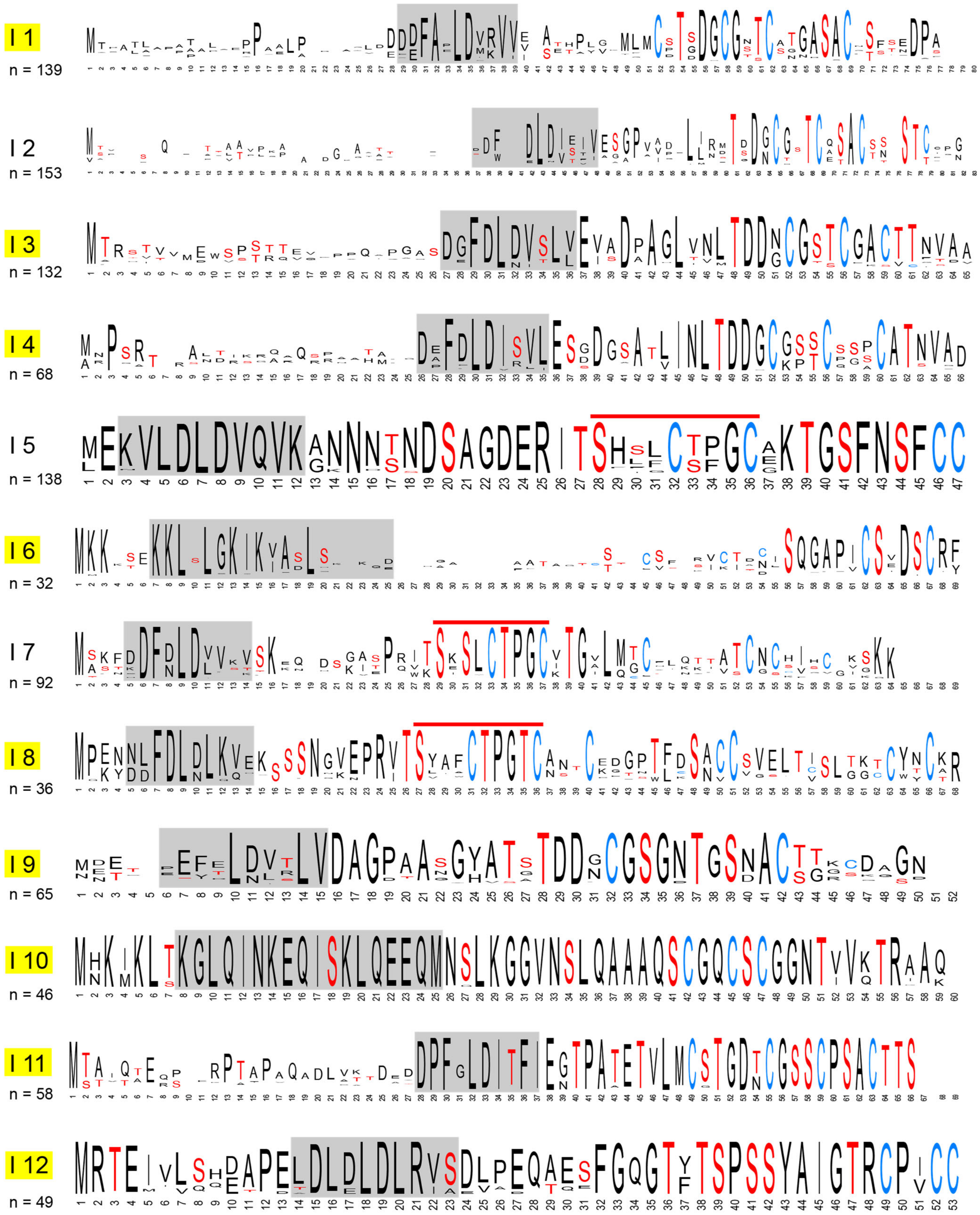

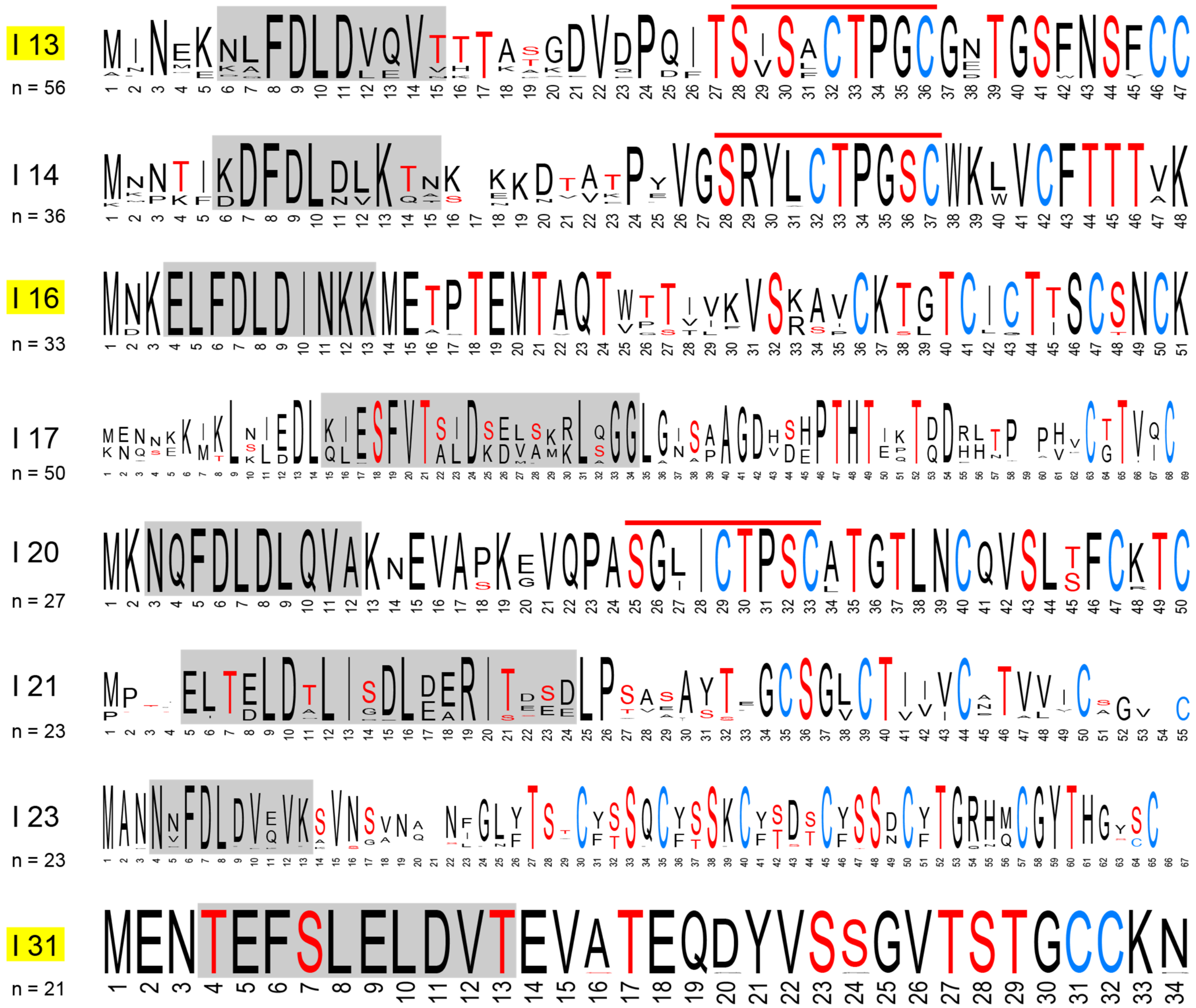
I 13
n = 56
I 14
n = 36
I 16
n = 33
I 17
n = 50
I 20
n = 27
I 21
n = 23
I 23
n = 23
I 31
n = 21

**Supplementary Figure S5.** Sequence logos generated from alignments of class II precursor peptides in clusters with 20 or more members (in Figure 2) using WebLogo [10]. Conserved leader motifs that are shared among multiple clusters (from Figure S2) are boxed in gray and potential lipid II-binding motifs are overlined in red. Families with no previously characterized members are highlighted in yellow.

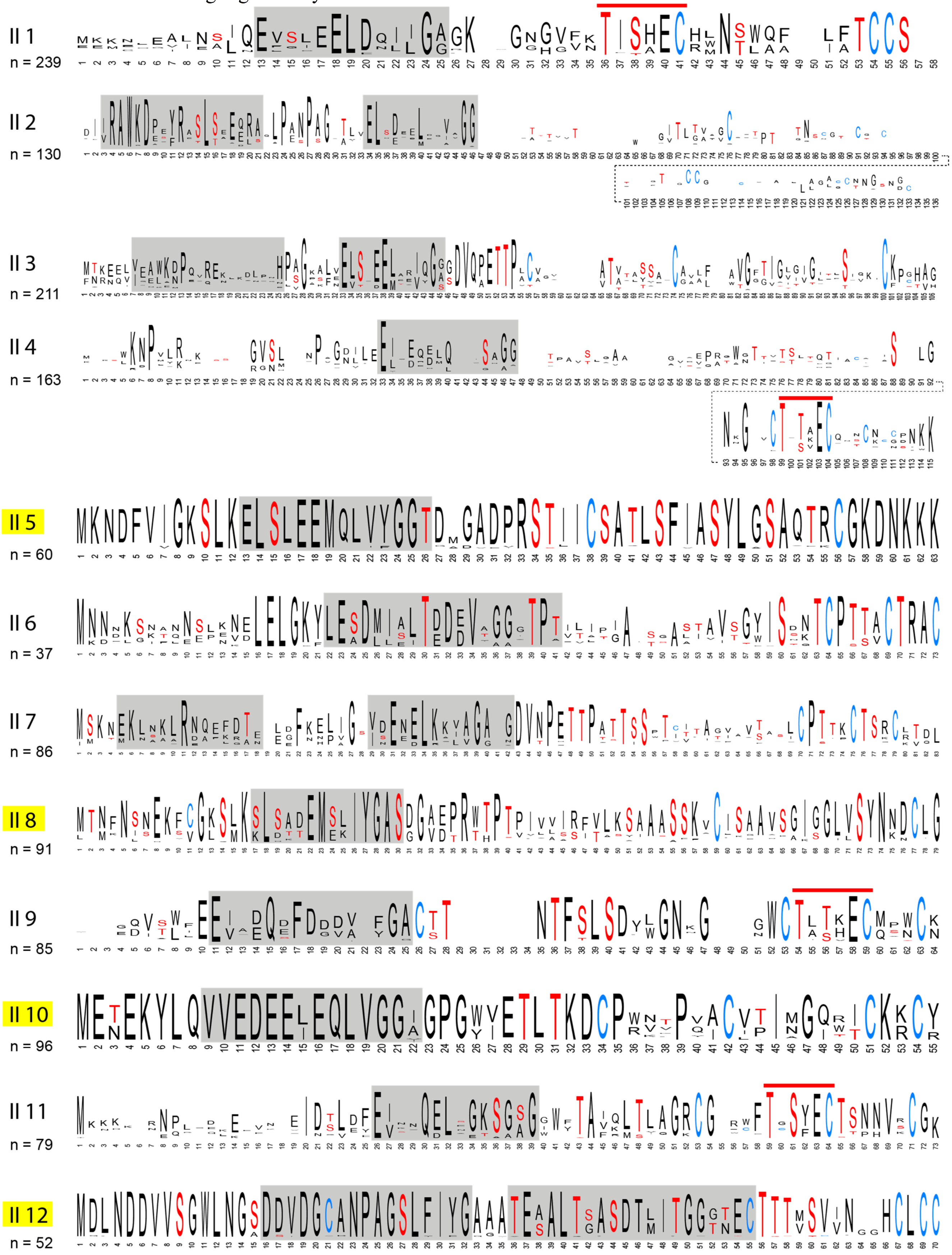

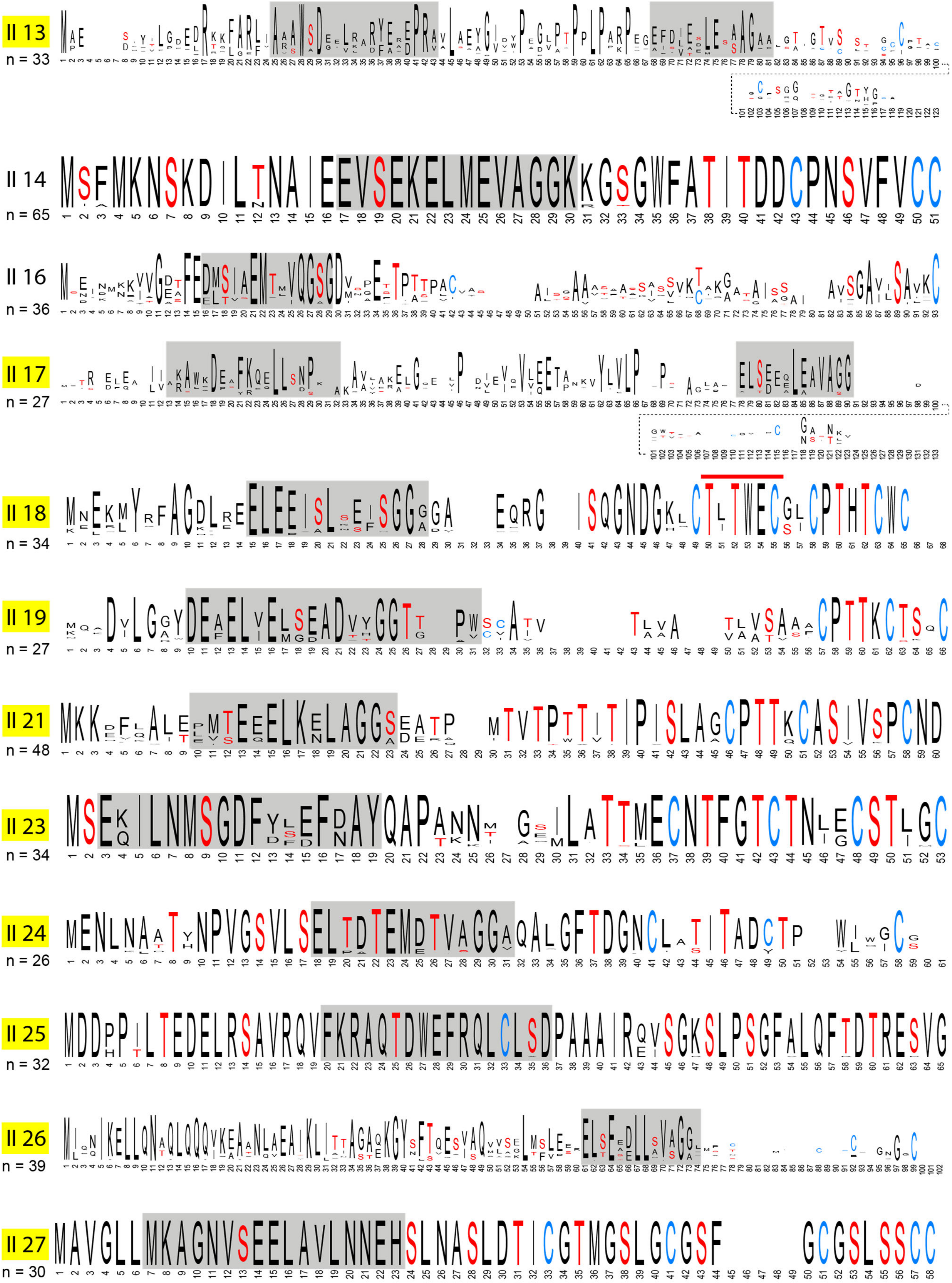

II 13
n = 33
II 14
n = 65
II 16
n = 36
II 17
n = 27
II 18
n = 34
II 19
n = 27
II 21
n = 48
II 23
n = 34
II 24
n = 26
II 25
n = 32
II 26
n = 39
II 27
n = 30

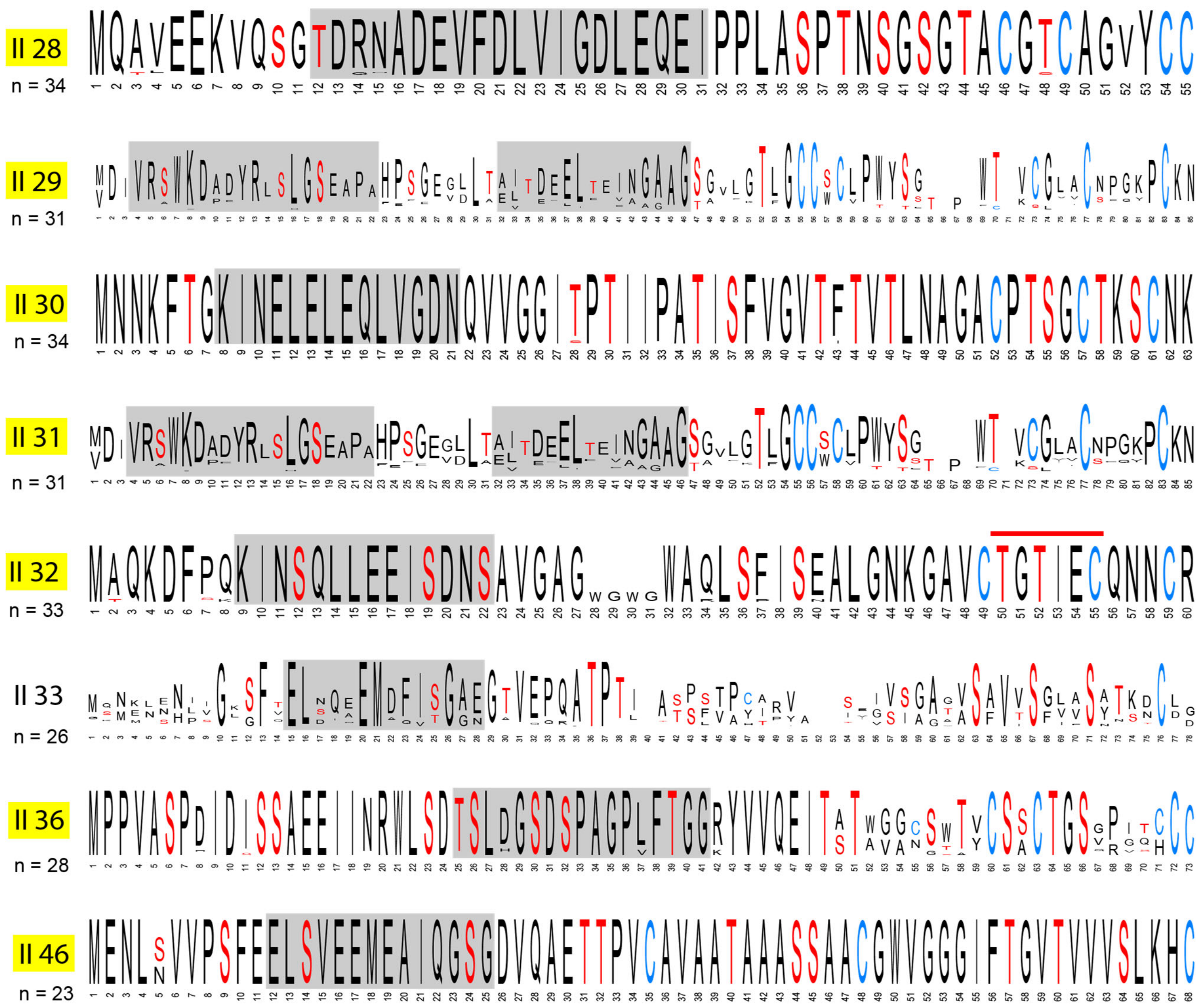

II 28
n = 34
II 29
n = 31
II 30
n = 34
II 31
n = 31
II 32
n = 33
II 33
n = 26
II 36
n = 28
II 46
n = 23

**Supplementary Figure S6.** Sequence logos generated from alignments of class III precursor peptides in clusters with 20 or more members (in Figure 2) using WebLogo [10]. Conserved leader motifs that are shared among multiple clusters (from Figure S2) are boxed in gray. Families with no characterized members are highlighted in yellow.

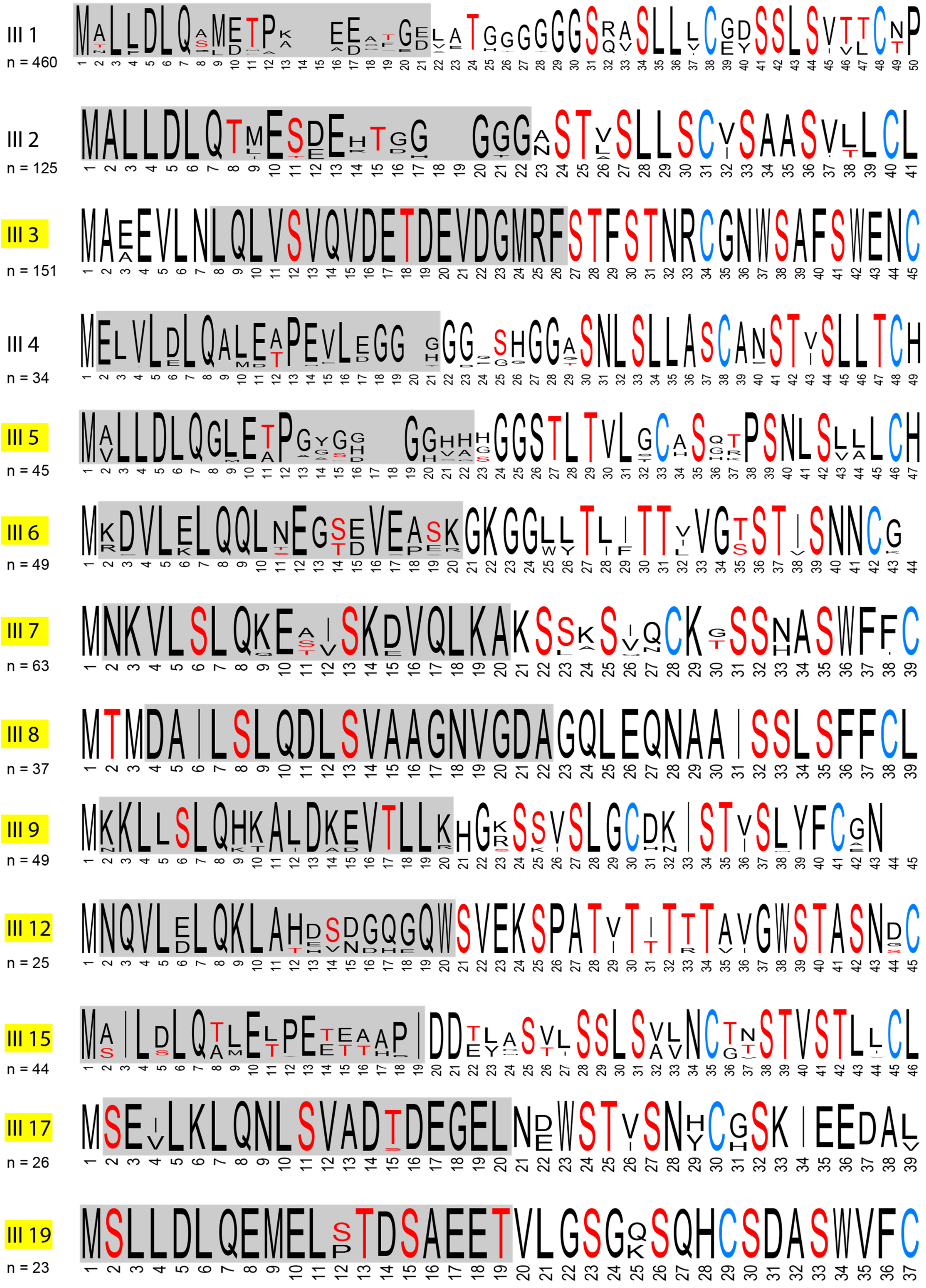

**Supplementary Figure S7.** Sequence logos generated from alignments of class IV precursor peptides in clusters with 20 or more members (in Figure 2) using WebLogo [10]. Conserved leader motifs that are shared among multiple clusters (from Figure S2) are boxed in gray. Families with no previously characterized members are highlighted in yellow.

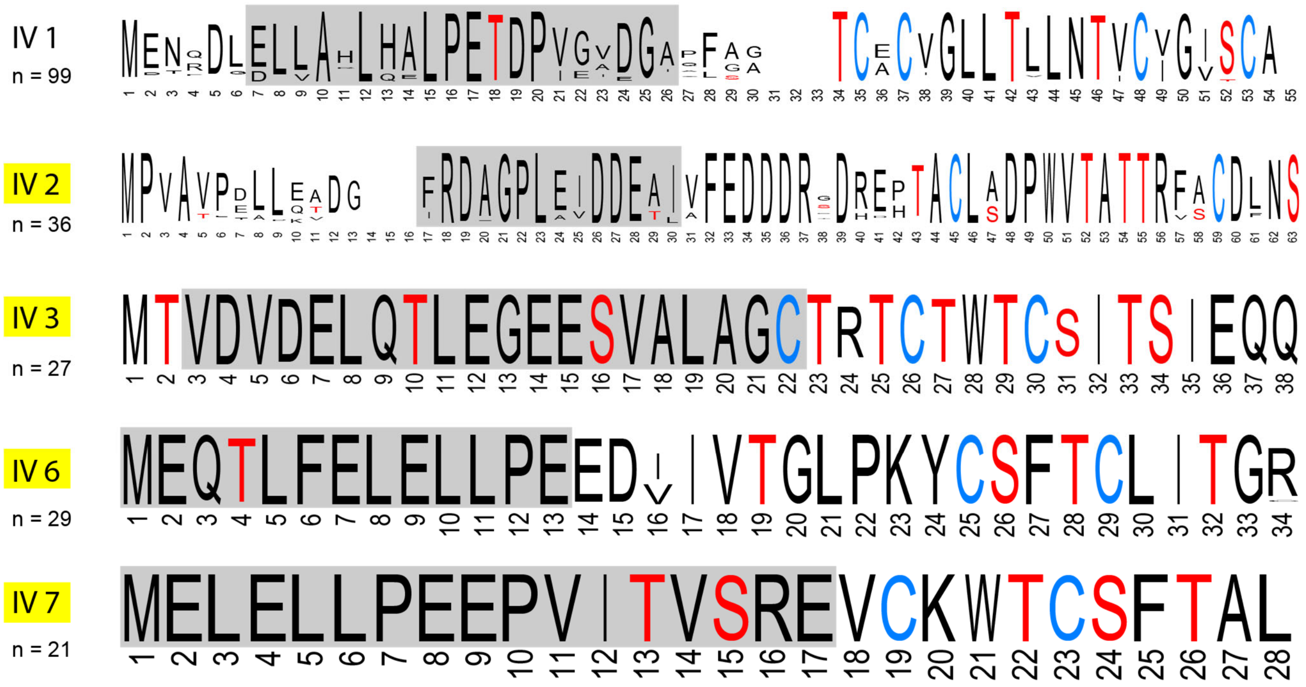

**Supplementary Table S6.** Twenty most abundant proteins in class I BGCs that belong to at least one Pfam family. If a protein has multiple domains from different Pfam families, those families are separated by a slash. Potential secondary modification enzymes are highlighted in yellow. Split LanBs, with the glutamylation and elimination domains on separate polypeptides, are denoted sLanB.

| Pfam families | Description | Count |
|---|---|---|
| LANC_like (PF05147) | LanC | 2,204 |
| Lant_dehydr_N (PF04738)/Lant_dehydr_C (PF14028) | LanB | 1,745 |
| ABC_tran (PF00005) | LanT | 657 |
| Lant_dehydr_C (PF14028) | LanB elimination domain | 562 (193 sLanBs) |
| PCMT (PF01135) | Protein-L-isoaspartate(D-aspartate) *O*-methyltransferase | 571 |
| Gallidermin (PF02052) | Gallidermin- and nisin-like precursor peptides | 426 |
| ABC2_membrane_4 (PF12730) | LanT | 363 |
| HTH_31 (PF13560) | transcriptional regulator | 307 |
| Lant_dehydr_N (PF04738) | PEARL | 304 (193 sLanBs) |
| ABC_membrane (PF00664)/ABC_tran (PF00005) | LanT | 283 |
| Lant_dehydr_C (PF14028)/PCMT (PF01135) | LanB elimination domain and Protein-L-isoaspartate(D-aspartate) *O*-methyltransferase fusion protein | 261 |
| Peptidase_C39 (PF03412)/ABC_membrane (PF12730)/ABC_tran (PF00005) | $LanT_P$ | 258 |
| HATPase_c_2 (PF13581) | Histidine kinase-like ATPase | 197 |
| Response_reg (PF00072)/Trans_reg_C (PF00486) | transcriptional regulator | 180 |
| Peptidase_S8 (PF00082) | $LanP_A$ | 172 |
| Flavoprotein (PF02441) | LanD | 146 |
| Acetyltransf_1 (PF00583) | *N*-acetyltransferase | 142 |
| DUF397 (PF04149) | Domain of Unknown Function | 140 |
| MFS_1 (PF07690) | Major Facilitator Superfamily protein | 128 |
| Leukocidin (PF07968) | Leukocidin/Hemolysin toxin family protein | 127 |

**Supplementary Table S7.** Twenty most abundant proteins in class II BGCs that belong to at least one Pfam family. If a protein has multiple domains from different Pfam families, those families are separated by a slash. Potential tailoring enzymes are highlighted in yellow.

| Pfam families | Description | Count |
|---|---|---|
| DUF4135 (PF13575)/LANC_like (PF05147) | LanM | 2,163 |
| Peptidase_C39 (PF03412)/ABC_membrane (PF12730)/ABC_tran (PF00005) | $LanT_P$ | 1,270 |
| ABC_tran (PF00005) | LanT | 886 |
| ABC2_membrane_4 (PF12730) | LanT | 761 |
| Mersacidin (PF16934) | Mersacidin-like precursor peptide | 705 |
| L_biotic_typeA (PF04604) | Type A lanthipeptide precursor peptide | 532 |
| HTH_3 (PF01381) | transcriptional regulator | 348 |
| Lantibiotic_a (PF14867) | Alpha precursor peptide | 339 |
| Response_reg (PF00072)/GerE (PF00196) | transcriptional regulator | 308 |
| Peptidase_S8 (PF00082) | $LanP_A$ | 291 |
| ABC_membrane (PF00664)/ABC_tran (PF00005) | LanT | 222 |
| ABC_tran (PF00005)/DUF4162 (PF13732) | LanT | 213 |
| FMN_red (PF03358) | Flavin mononucleotide reductase | 186 |
| HisKA (PF00512)/HATPase_c (PF02518) | 2-component response regulator | 180 |
| Response_reg (PF00072)/Trans_reg_C (PF00486) | transcriptional regulator | 164 |
| GerE (PF00196) | transcriptional regulator | 140 |
| Nhase_alpha (PF02979) | precursor peptide with nitrile hydratase family leader peptide | 119 |
| Nif11 (PF07862) | precursor peptide with Nif11 family leader peptide | 109 |
| FtsX (PF02687) | FtsX-like permease family | 104 |
| LANC_like (PF05147) | LanC (split LanM possibly from sequencing errors) | 95 |

**Supplementary Table S8.** Twenty most abundant proteins in class III BGCs that belong to at least one Pfam family. If a protein has multiple domains from different Pfam families, those families are separated by a back slash. Potential tailoring enzymes are highlighted. $LanP_P$ is a Pro oligopeptidase that is distinct from the $LanP_A$ subtilin-like S8 peptidases.

| Pfam families | Description | Count |
|---|---|---|
| ABC_membrane (PF00664)/ABC_tran (PF00005) | LanT | 1,514 |
| Pkinase (PF00069)/LANC_like (PF05147) | LanKC | 1,511 |
| ABC_tran (PF00005) | LanT | 781 |
| Response_reg (PF00072)/GerE (PF00196) | transcriptional regulator | 715 |
| GerE (PF00196) | transcriptional regulator | 263 |
| MFS_1 (PF07690) | Major facilitator superfamily protein | 243 |
| adh_short (PF00106) | short chain dehydrogenase | 156 |
| GAF_2 (PF13185)/PAS_3 (PF08447)/GAF_2 (PF13185)/SpoIIE (PF07228)/HATPase_c_2 (PF13581) | Unknown | 135 |
| trypsin (PF00089) | Protease | 133 |
| Pkinase (PF00069) | protein kinase | 133 |
| HTH_20 (PF12840) | transcriptional regulator | 123 |
| Acetyltransf_1 (PF00583) | N-acetyltransferase | 122 |
| BPD_transp_1 (PF00528) | Binding-protein-dependent transport system, inner membrane component | 121 |
| FtsX (PF02687)/FtsX (PF02687) | FtsX-like permease | 119 |
| GAF_2 (PF13185)/PAS_4 (PF08448)/GAF_2 (PF13185)/SpoIIE (PF07228) | Unknown | 114 |
| Acetyltransf_3 (PF13302) | *N*-acetyltransferase | 113 |
| Peptidase_S9 (PF00326) | $LanP_P$ | 112 |
| FecCD (PF01032) | FecCD transport family | 102 |
| Mac (PF12464) /Hexapep (PF00132) | Acetyltransferase | 97 |
| DUF4265 (PF14085) | Domain of unknown function | 93 |

**Supplementary Table S9.** Twenty most abundant proteins in class IV BGCs that belong to at least one Pfam family. If a protein has multiple domains from different Pfam families, those families are separated by a back slash. Known class IV BGCs comprise only 4 genes, so these entries may include proteins encoded by genes that are not part of the gene cluster. Potential tailoring enzymes are highlighted.

| Pfam families | Description | Count |
|---|---|---|
| Pkinase (PF00069)/LANC_like (PF05147) | LanL | 340 |
| ABC_membrane (PF00664)/ABC_tran (PF00005) | LanT | 164 |
| Peptidase_S9 (PF00326) | $LanP_P$ | 112 |
| MFS_1 (PF07690) | Major facilitator Superfamily protein | 101 |
| ABC2_membrane (PF01061) | LanT | 83 |
| ABC_tran (PF00005)/DUF4162 (PF13732) | LanT | 82 |
| HATPase_c_2 (PF13581) | Histidine kinase-like ATPase domain | 48 |
| ABC_tran (PF00005) | LanT | 48 |
| Nif3 (PF01784) | NGG1p interacting factor 3 | 46 |
| NAD_binding_10 (PF13460) | NAD(P)H-binding protein | 35 |
| STAS_2 (PF13466) | STAS domain containing protein | 31 |
| DAO (PF01266) | FAD dependent oxidoreductase | 30 |
| BPD_transp_1 (PF00528) | Binding-protein-dependent transport system, inner membrane component | 30 |
| TrmK (PF04816) | N-methyltransferase | 29 |
| Glycos_transf_2 (PF00535) | Glycosyl transferase | 28 |
| SBP_bac_3 (PF00497) | Bacterial extracellular solute-binding protein | 26 |
| Methyltransf_19 (PF04672) | Methyltransferase | 26 |
| GDP_Man_Dehyd (PF16363) | GDP-mannose 4,6-dehydratase | 26 |
| SNF2_assoc (PF08455)/SNF2_N (PF00176)/Helicase_C (PF00271) | Helicase | 24 |
| Response_reg (PF00072)/Sigma70_r4_2 (PF08281) | Response regulator | 25 |

**Supplementary Table S10.** Distribution of Pfams that are in the 20 most abundant protein families in one class among the other three classes. Values are presented on the basis of domains, so Pfam families that occur in multidomain proteins and single domain proteins are counted together. The Pfam are listed in order of decreasing frequency.

| Pfam protein family | Top 20 most abundant in class | Class I | Class II | Class III | Class IV |
|---|---|---|---|---|---|
| PCMT (PF01135) | I | 835 | 1 | 1 | 0 |
| Flavoprotein (PF02441) | I | 146 | 26 | 24 | 4 |
| Acetyltransf_1 (PF00583) | I, III | 149 | 47 | 142 | 12 |
| FMN_red (PF03358) | II | 34 | 186 | 7 | 4 |
| adh_short (PF00106) | III | 57 | 30 | 157 | 19 |
| Acetyltransf_3 (PF13302) | III | 57 | 12 | 113 | 9 |
| Mac (PF12464) | III | 0 | 0 | 97 | 0 |
| TrmK (PF04816) | IV | 0 | 0 | 0 | 29 |
| DAO (PF01266) | IV | 6 | 1 | 9 | 30 |
| NAD_binding_10 (PF13460) | IV | 21 | 10 | 19 | 38 |
| GDP_Man_Dehyd (PF16363) | IV | 2 | 0 | 65 | 26 |
| Glycos_transf_2 (PF00535) | IV | 36 | 8 | 39 | 29 |
| Methyltransf_19 (PF04672) | IV | 64 | 12 | 9 | 26 |

**Supplementary Figure S8.** Example putative biosynthetic gene clusters encoding the enzymes in Table S10. LanA genes that were not annotated in the genome are indicated in red. Because BGC boundaries are not known, the genes encoding the noted enzymes may or may not be part of the lanthipeptide BGC for all panels of Supplementary Figure S8.

### *O*-methyltransferase containing clusters

#### Class I

*Streptomyces clavuligerus* ATCC 27064

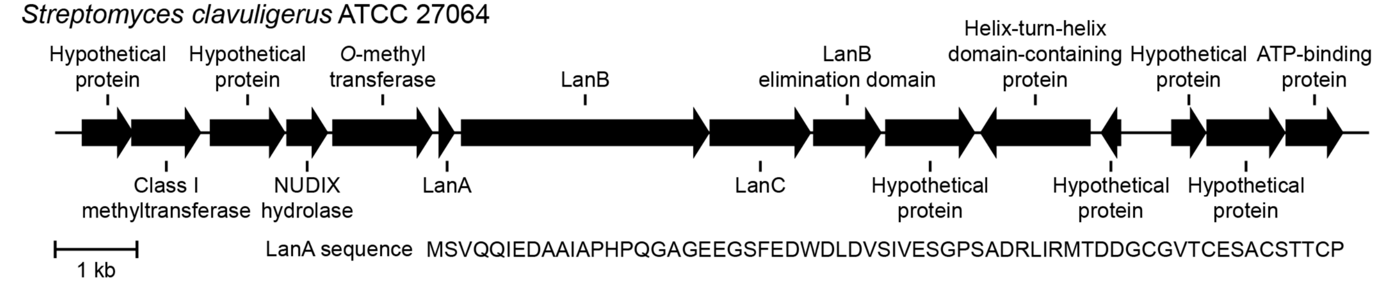


#### Class II

*Marinicella sediminis* strain F2

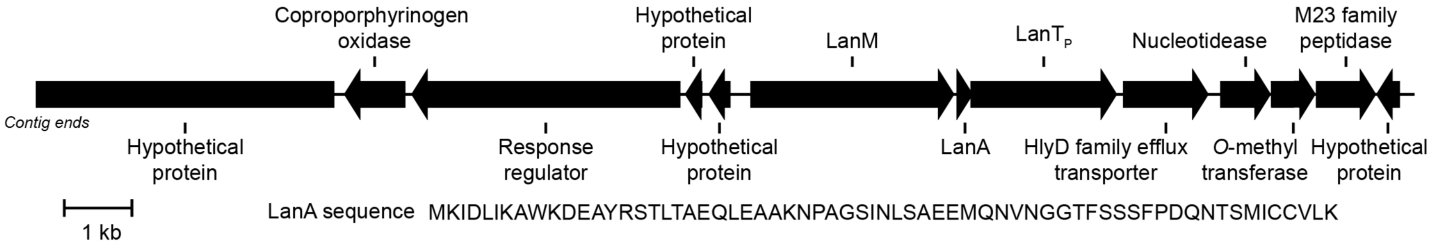


#### Class III

*Streptomyces varsoviensis* strain NRRL ISP-5346

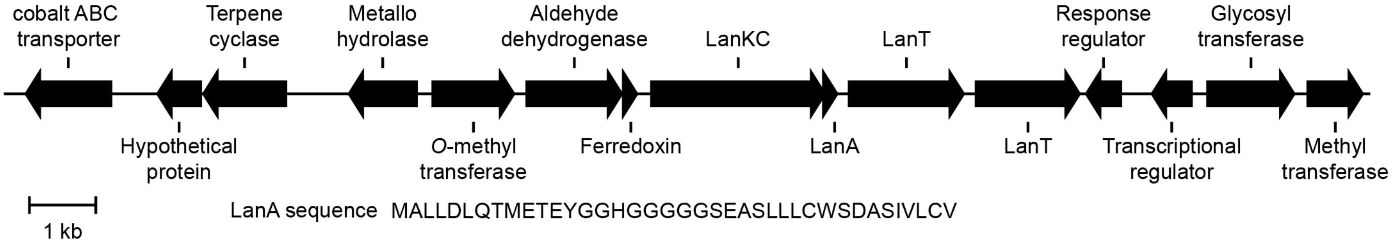

**Supplementary Figure S8 continued.**

### Flavoprotein containing clusters

#### Class I

*Brevibacillus* sp. BC25

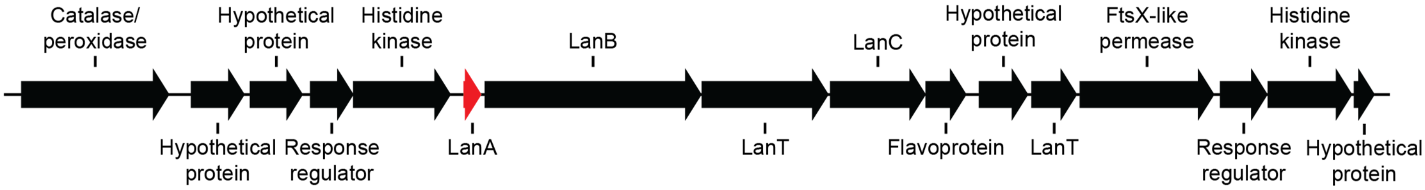


1 kb

LanA sequence MQKDLFDLDVQVKEVNQVQSDSVVSDIICTTFCSVTWCQSNCC

#### Class II

*Bacillus velezensis* strain GH1-13

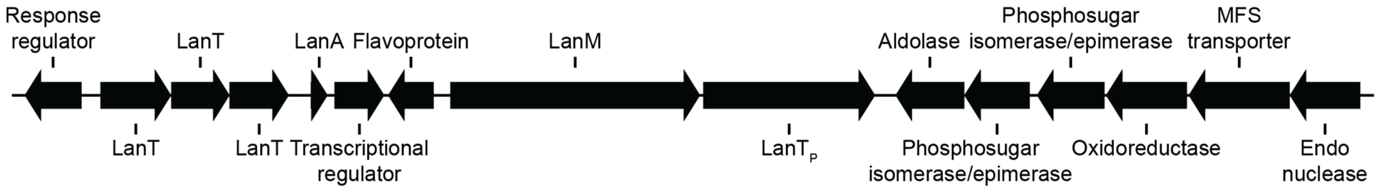


1 kb

LanA sequence MSQEAIIRSWKDPFSRENSTQNPAGNPFSELKEAQMDKLVGAGDMEAACTFTLPGGGGVCTLTSECIC

#### Class III

*Streptomyces bicolor* strain NRRL B-5348

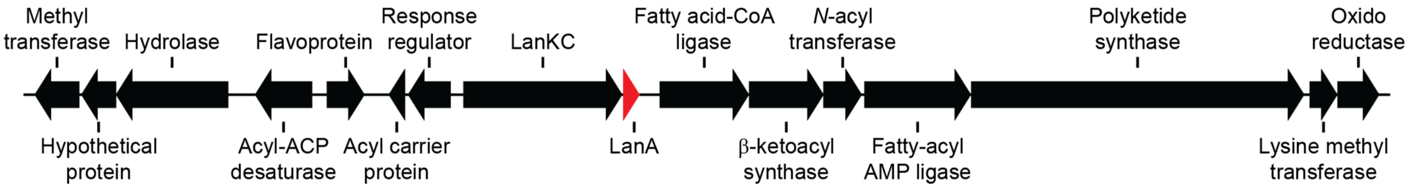


1 kb

LanA sequence MTESVLDLQELETSEEETALMAASASSWNC

#### Class IV

*Streptomyces* sp. AcH 505

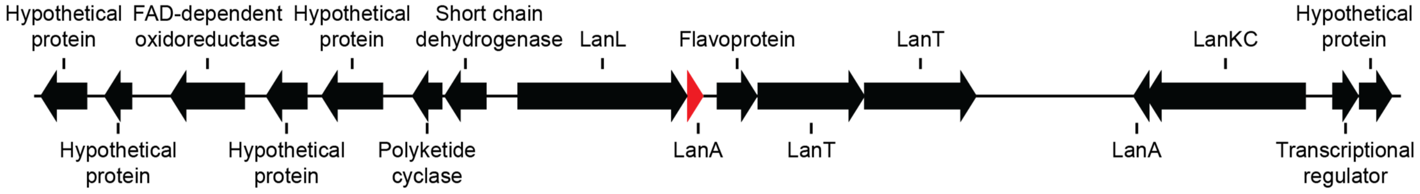


1 kb

LanL-LanA sequence MESDLDALQLLTGEEAQEQSLILCEGHTCNGGSTCAISCNVTN

LanKC-LanA sequence MQQVLNLQELETEHTVDENGVIIASDFSTFACDSNVSVVVC

**Supplementary Figure S8 continued.**

### Acyltransferase (Acyltransf_1) containing clusters

#### Class I

*Paenibacillus polymyxa* strain NCTC10343

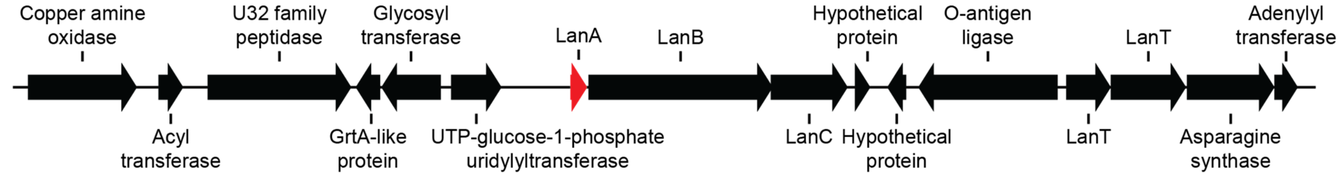


1 kb

LanA sequence MNSPELVFFEQEDTLDLDLQINDLTLKQAKNPCTSTVTCSVSRCLGTHVTCECWC

#### Class II

*Thermoanaerobaculum aquaticum* strain MP-01

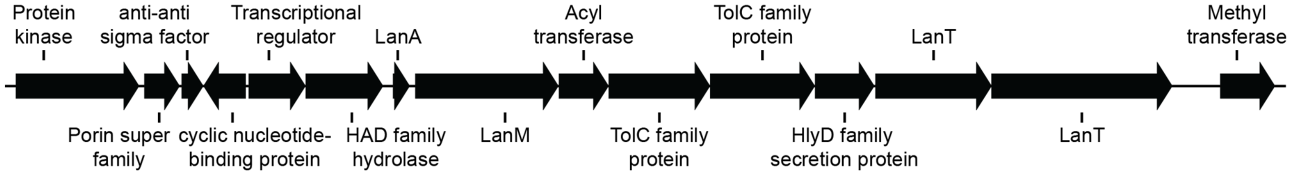


1 kb

LanA sequence MPEAVTLTRGEVNDLIAGFATKNPEYRKALLAKPREVVGAQLGTSIPPSVQVKVIEEKPNEFYVIIPHVA KEGQELSDADLEQVAGGKKDTYKCEVQGLGFGTRVEINASLF

#### Class III

*Streptomyces* sp. CNS654

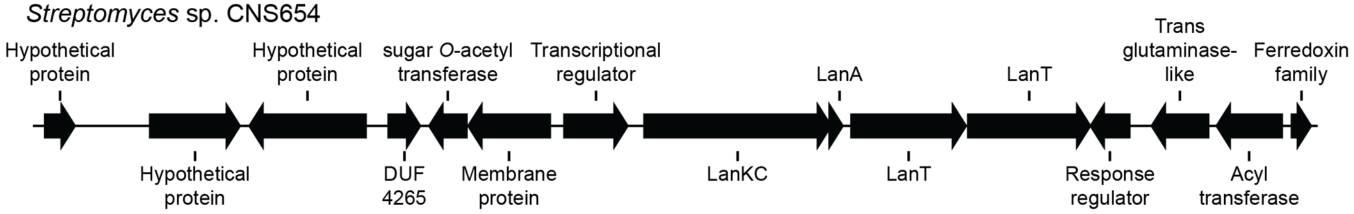


1 kb

LanA sequence MALLDLQAMDTPAEDSFGELATGSQVSLLVCEYSSLSVVLCTP

#### Class IV

*Streptomyces baarnensis* strain NRRL B-2842

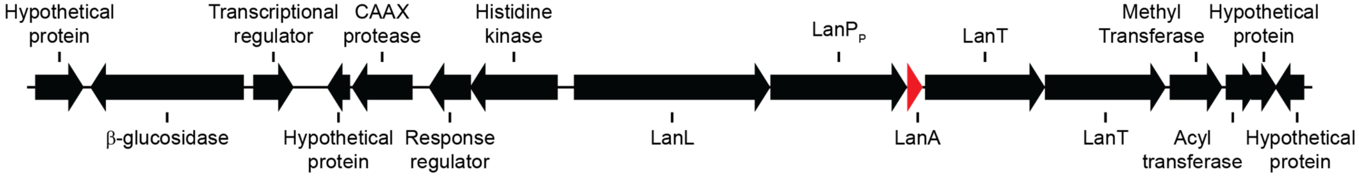


1 kb

LanA sequence MIDNTMTVDVDELQTLEGEESVALAGCTRTCTWTCSITSIEQQ

**Supplementary Figure S8 continued.**

### FMN reductase (FMN_red) containing clusters

#### Class I

*Listeria monocytogenes* strain AL-OM-1-WH2

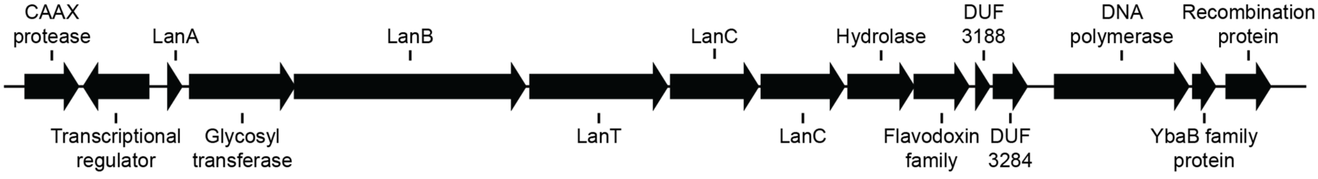


LanA sequence MSGEKDFDLNAQLKSENNTEVTAASWWMFTIITISKAVSGTYSSSETAKNCSMSSCKKC

#### Class II

*Bacillus cereus* VD166

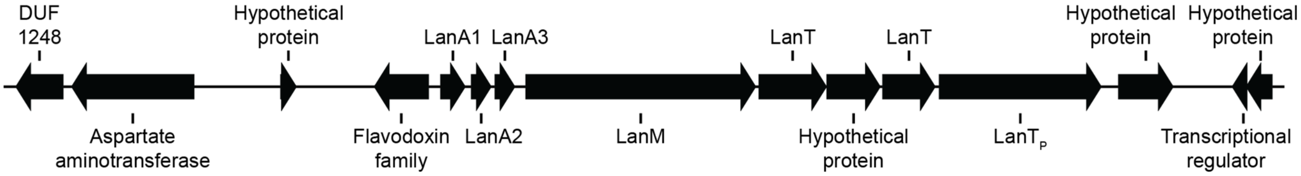


LanA1 sequence MIVTVVIVTKKNNLRRNFIMIKSNLLKDPVLKKKLAINLDNPIGDIGEEILEQDVNGQVGGLSPAAISLA GTVISVVSVAGGGIIKAASAKAQNPGRYCTISAECFAGERCD

LanA2 sequence MFSQKSYLLKNPVAKNKFGASMTNPSGDIITEIQEQELEAVNGGITPTVGWSAVAVSTLTVNSVLVVSVS ATNPGRVCTISAECSSGYKRCD

LanA3 sequence MHMNHKLMKNPVAKNKLKVLDESPAGDLLEEIQEQELQSNGYGGISPSVVATISVVSAVSIAGTSVVSAS ATNPGRYCTWSAECSWSQKRCD

#### Class III

*Streptomyces* sp. CNS654

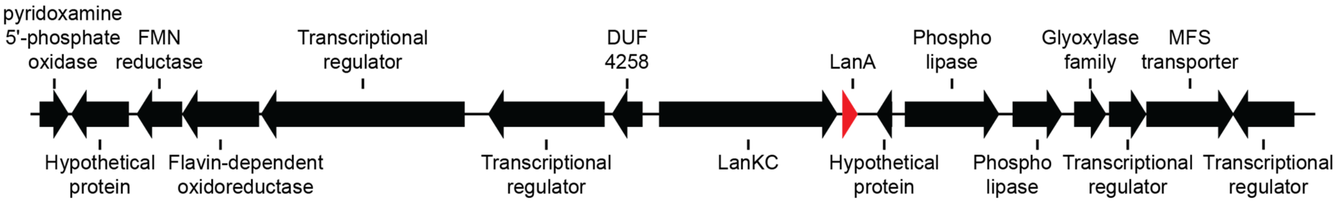


LanA sequence MSILNLQQLEIAEEEGGLLLMSSASFQCHNTSFH

#### Class IV

*Streptomyces misionensis* strain DSM 40306

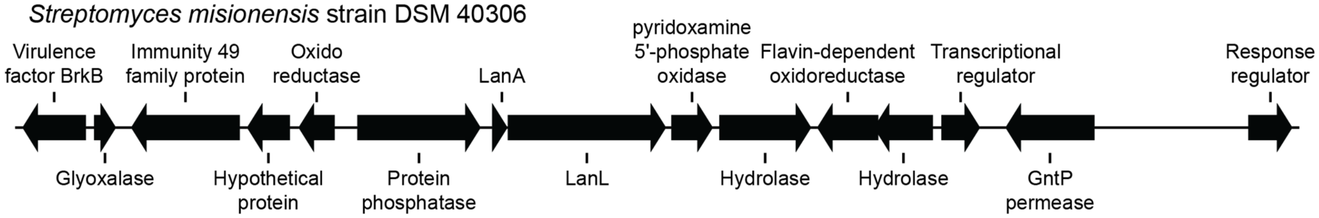


LanA sequence MPVAVPELLEADGFRDAGPLEIDDEAIVFEDDDRGDREHTACLSDPWVTATTRFSCDLNS

**Supplementary Figure S8 continued.**

### Short chain dehydrogenase (adh_short) containing clusters

#### Class I

*Frankia* sp. QA3

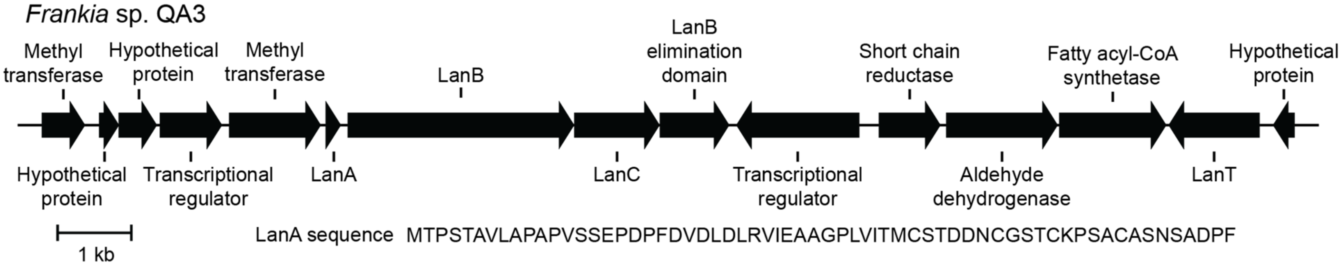


LanA sequence MTPSTAVLAPAPVSSEPDPFDVDLDLRVIEAAGPLVITMCSTDDNCGSTCKPSACASNSADPF

#### Class II

*Bacillus thuringiensis* MC28

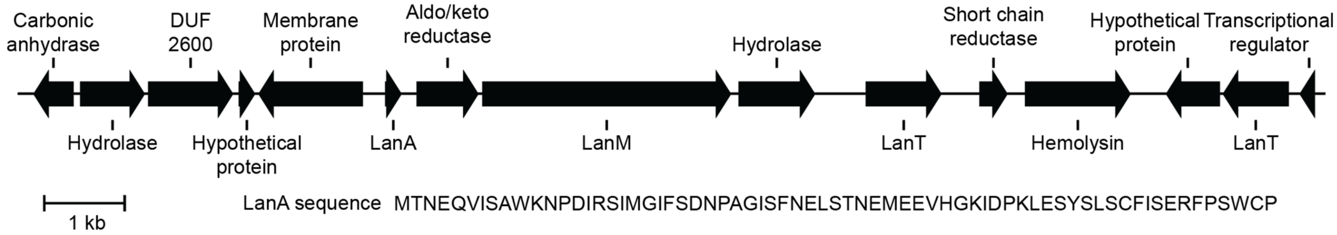


LanA sequence MTNEQVISAWKNPDIRSIMGIFSDNPAGISFNELSTNEMEEVHGKIDPKLESYSLSCFISERFPSWCP

#### Class III

*Rhodococcus erythropolis* SK121

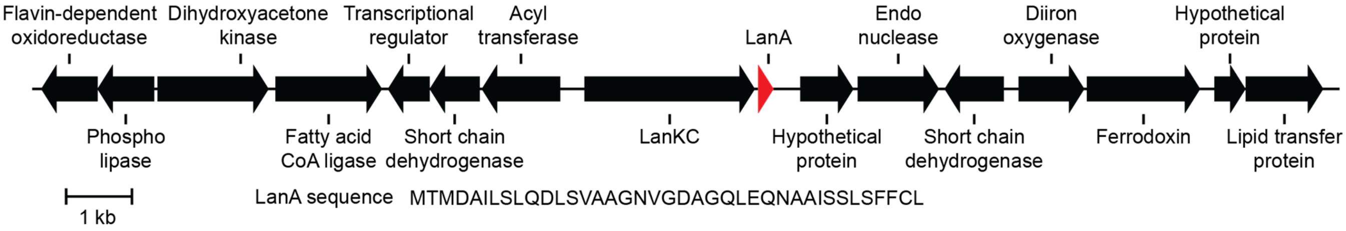


LanA sequence MTMDAILSLQDLSVAAGNVGDAGQLEQNAAISSLSFFCL

#### Class IV

*Streptomyces flavochromogenes* strain NRRL B-2684

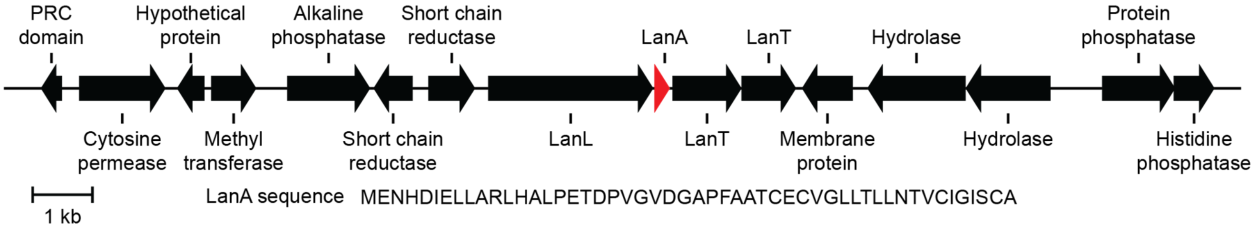


LanA sequence MENHDIELLARLHALPETDPVGVDGAPFAATCECVGLLTLLNTVCIGISCA

**Supplementary Figure S8 continued.**

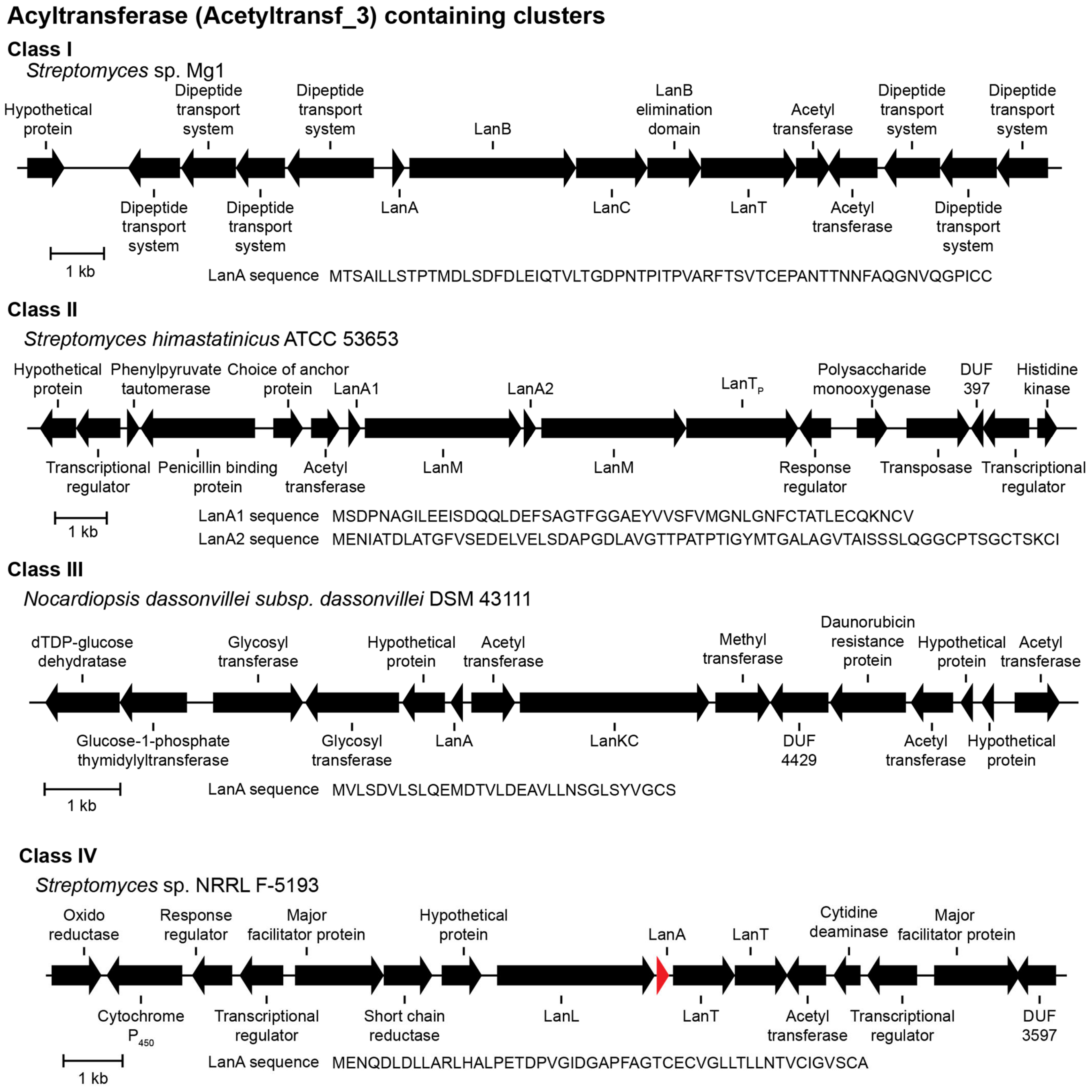

**Supplementary Figure S8 continued.**

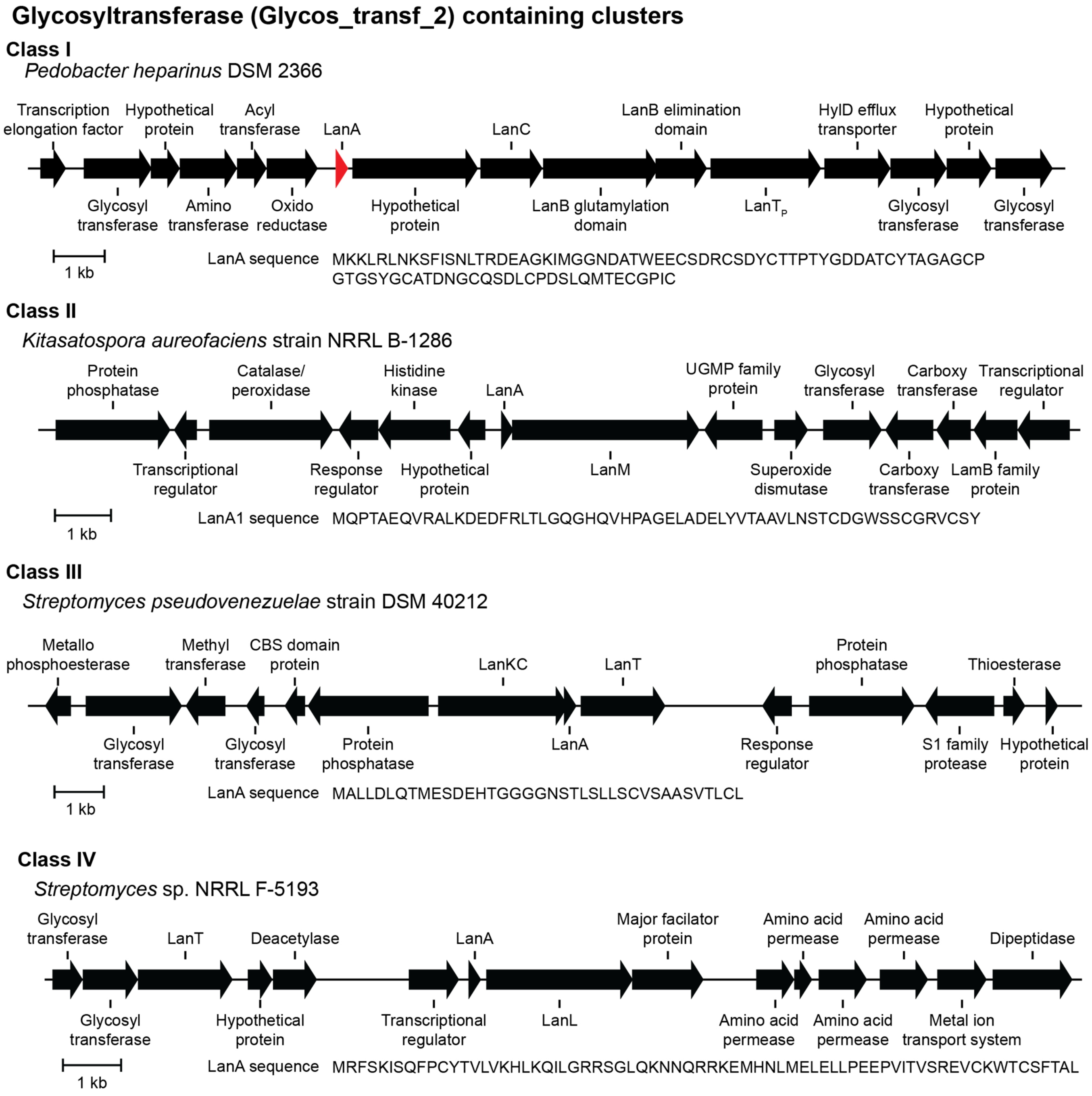

**Supplementary Figure S8 continued.**

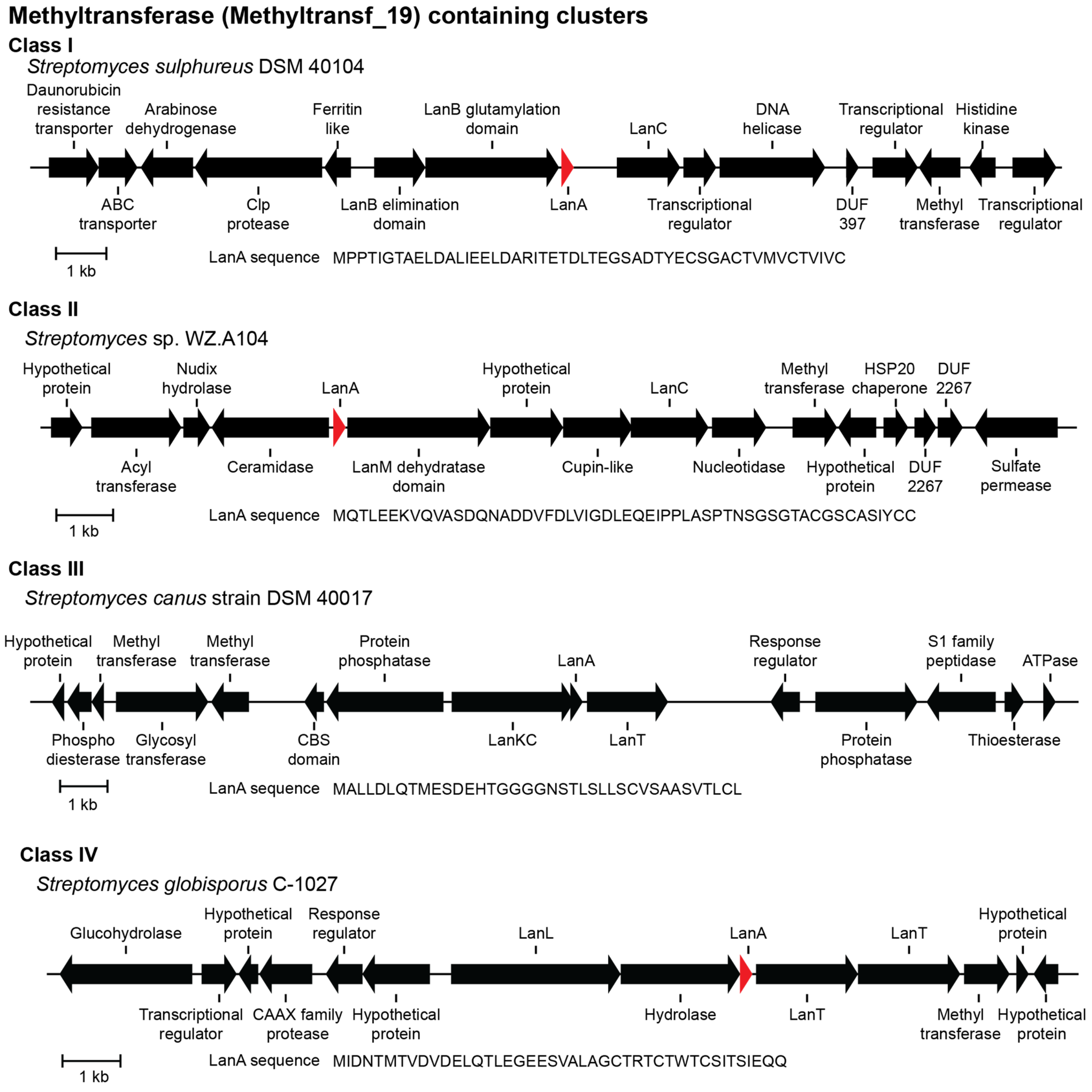

**Supplementary Table S11.** Distribution of select Pfam protein families from BGCs. The enzymes belonging to these families potentially carry-out secondary post-translational modifications. Values are presented on the basis of domains, so Pfam protein families that occur in multidomain proteins or single domain proteins are counted together. NRPS: non-ribosomal peptide synthetase, PKS: polyketide synthase, FAS: fatty acid synthase.

| Pfam protein families | Class I | Class II | Class III | Class IV |
|---|---|---|---|---|
| YcaO (PF02624) | 4 | 42 | 2 | 0 |
| Radical_SAM (PF04055) | 39 | 49 | 42 | 43 |
| p450 (PF00067) | 45 | 44 | 50 | 23 |
| Condensation (PF00668) (NRPS) | 45 | 31 | 10 | 2 |
| Ketoacyl-synt_C (PF02801) (PKS/FAS) | 34 | 5 | 92 | 8 |
| ADH_N (PF08240) (zinc-dependent dehydrogenase) | 56 | 63 | 38 | 27 |
| 2OG-FeII_Oxy_3 (PF13640) | 0 | 5 | 4 | 0 |

**Supplementary Figure S9.** Example biosynthetic gene clusters encoding the enzymes in Table S11. LanA genes that were not annotated in the genome are indicated in red. Because BGC boundaries are not known, the noted enzymes may or may not be part of the lanthipeptide BGC for all panels of Supplementary Figure S9.

### YcaO containing clusters

#### Class I

*Actinokineospora enzanensis* DSM 44649

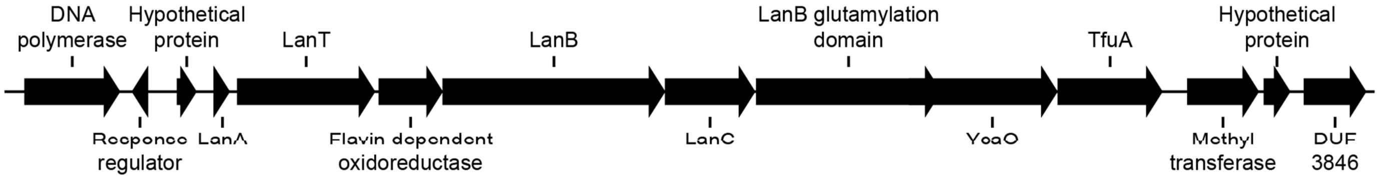


LanA sequence MTQMIAGPGFDLDLTVDEEIFSAPGGGSQATASTADNPDGTFTFTTTCWGTCAVTCGDCHTLECCTH

#### Class II

*Chryseobacterium vrystaatense* strain LMG 22846

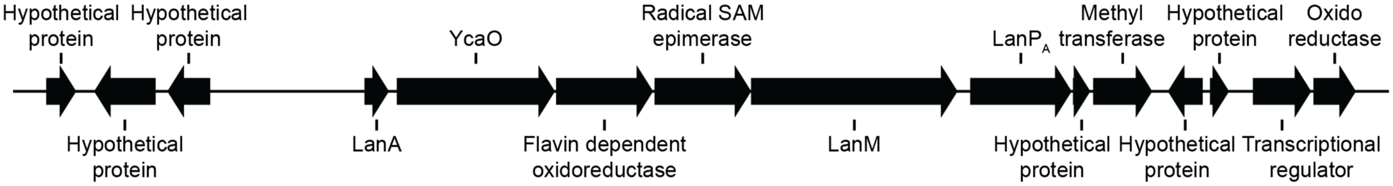


LanA sequence MENQAVRVISEVTAKAQANPDFAREYVSNPNRVLSDAGMQIPEGMNIHVIVGAPANSEIPNSTSTDVYLL LPQVNEEIKDESLATAAAASCQSTSSTCVTVPSCVSCVSSASTNSCS

#### Class III

*Nocardiopsis valliformis* DSM 45023

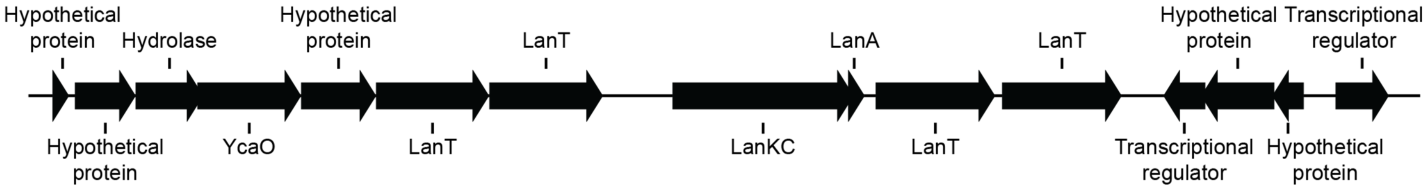


LanA sequence MTLLDLQSLETAKNEAHGEAGGTSTASLLLCGDSSLSITTC

**Supplementary Figure S9 continued.**

### Radical SAM containing clusters

#### Class I

*Tannerella forsythia* 92A2

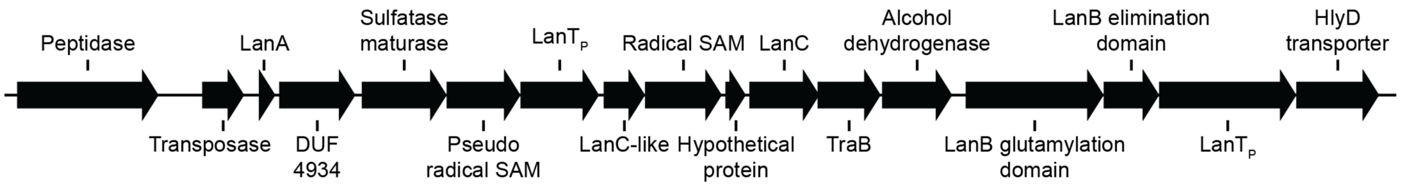


1 kb

LanA sequence MKKLKKIKLNQLCKAELNSREMAQLTGGLCCGCGCHGPSSTVDNQNANAENGYGSIGGNKICWCW DGGAWVKTETK

#### Class II

*Butyrivibrio proteoclasticus* strain P18

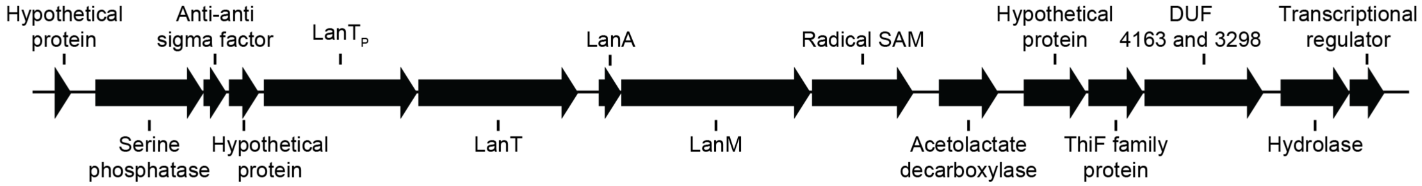


1 kb

LanA sequence MKTLQELHKEISANKDLQKAYAAAIENDAVLDFVKANGCDVTEDDIKEYISGIKKDDDTPLSVEDIENVS GGGLICKTVSVYKKMSKMAKKKGLIPDWYPC

#### Class III

*Kibdelosporangium aridum* strain A82846

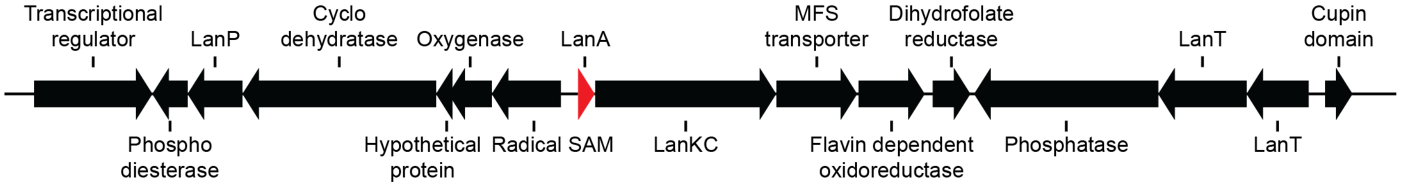


1 kb

LanA sequence MSDEKPQSSEEQSENEEVVAHAVLDLQKIQTTMPAGQIGNPAGSCSSCFAASCG

#### Class IV

*Kibdelosporangium aridum* strain A82846

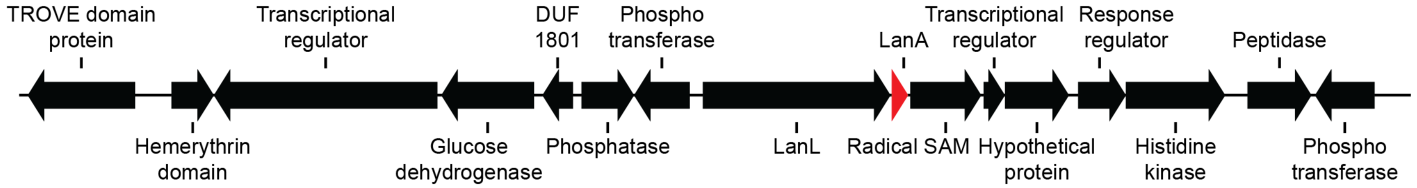


1 kb

LanA sequence MSDEKPQSSEEQSENEEVVAHAVLDLQKIQTTMPAGQIGNPAGSCSSCFAASCG

**Supplementary Figure S9 continued.**

### Cytochrome $P_{450}$ containing clusters

#### Class I

*Actinomadura madurae* strain DSM 43067

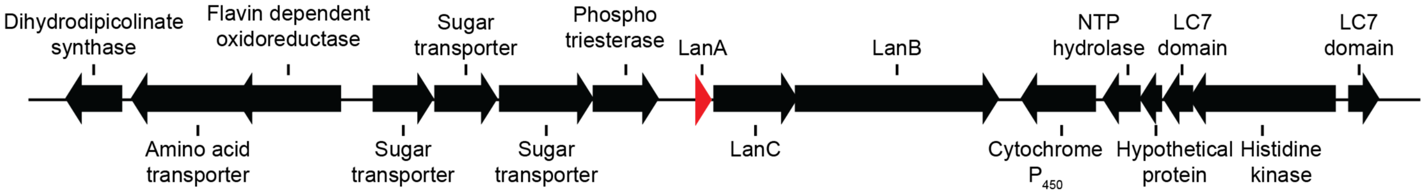


1 kb

LanA sequence MTISLTDQAGVDAQDEFDLDVRVSELSTSAEILDCTDSCPCFTAPCQSTPTCLEGI

#### Class II

*Lentzea terrae* strain NEAU-LZS 42

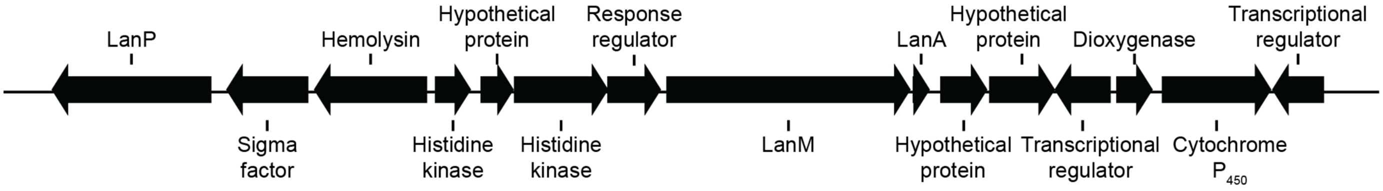


1 kb

LanA sequence MKNATIIRAWKDAAFRATLPAGAVPAHPAGSAVVKTELLGAAMGTTPICTDGGPRCTDSGPHCS

#### Class III

*Streptomyces* sp. NRRL F-4428

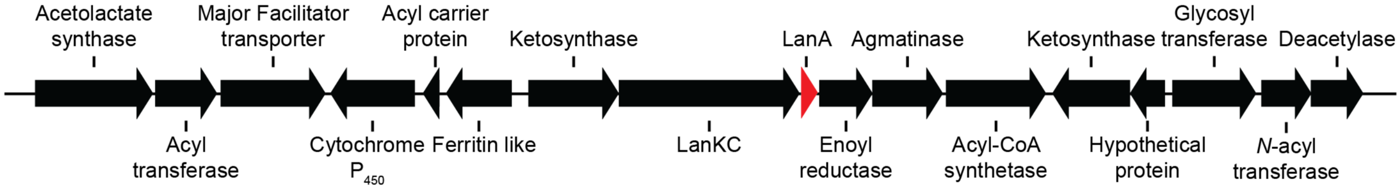


1 kb

LanA sequence MSIVLDLQGLEVPAEETTRVASTASNHC

#### Class IV

*Streptomyces* sp. NRRL F-5193

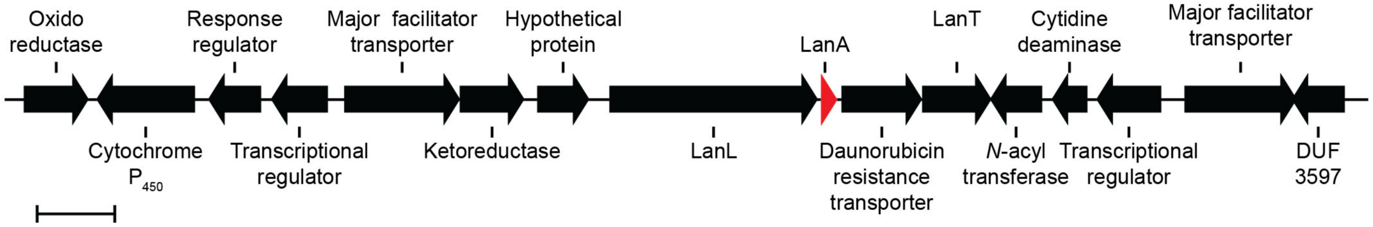


1 kb

LanA sequence MENQDLDLLARLHALPETDPVGIDGAPFAGTCECVGLLTLLNTVCIGVSCA

**Supplementary Figure S9 continued.**

### Nonribosomal peptide synthetase containing clusters

#### Class I

*Tumebacillus avium* strain AR23208

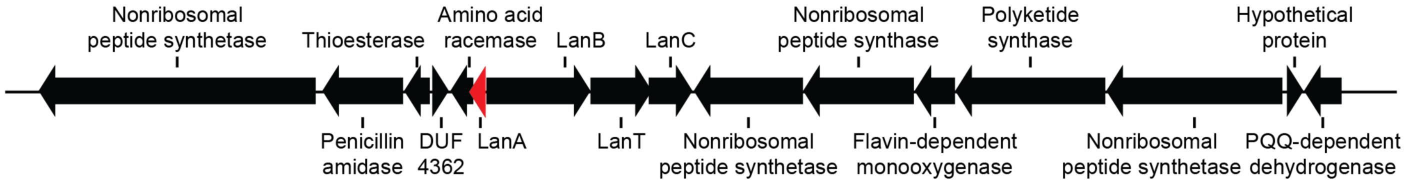


1 kb  LanA sequence MENHFDLDVQVGEATVDVTRRDLTGNTSCLISCDIDI

#### Class II

*Saccharothrix variisporea* strain DSM 43911

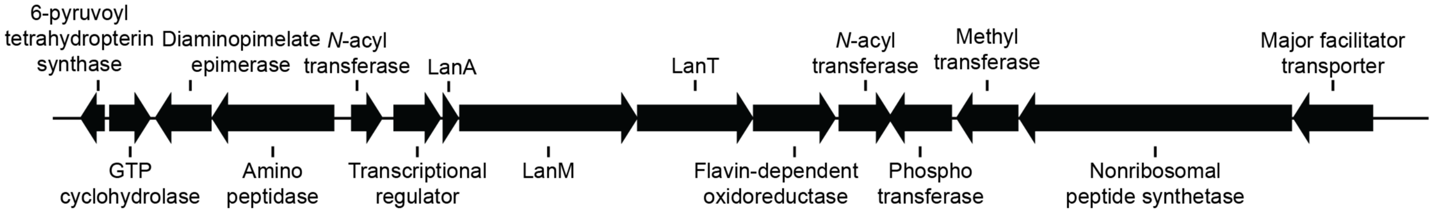


1 kb  LanA sequence MTTALLESAEELVARWRAGLDDVDNPAGPLFANGEFAEGDIVACNPPGSRCSSCTASRPVYCC

#### Class III

*Streptomyces* sp. NRRL F-4428

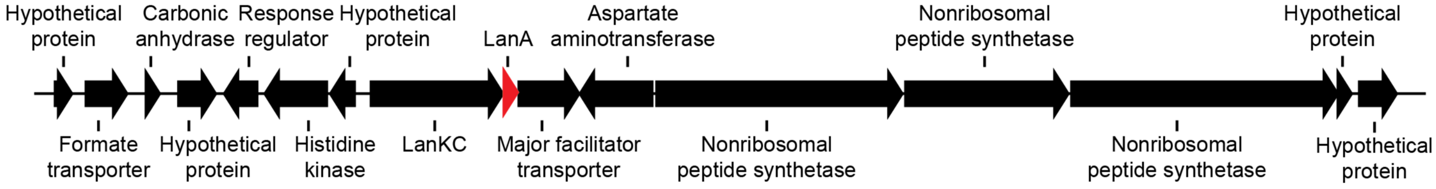


1 kb  LanA sequence MSNVLRLQALRPQTTNRQPVISSSYSFICCN

#### Class IV

*Streptomyces venezuelae*

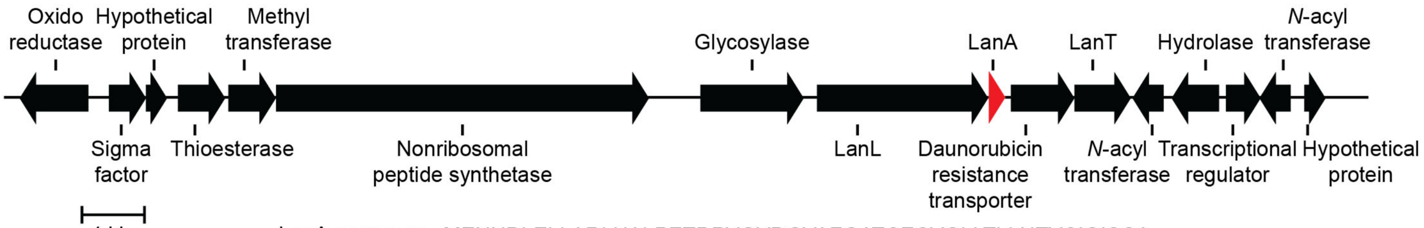


1 kb  LanA sequence MENNDLELLARLHALPETDPVGVDGVAFGATCECVGLLTLLNTVCIGISCA

**Supplementary Figure S9 continued.**

### Polyketide synthase containing clusters

#### Class I

*Streptomyces formicae* strain KY5

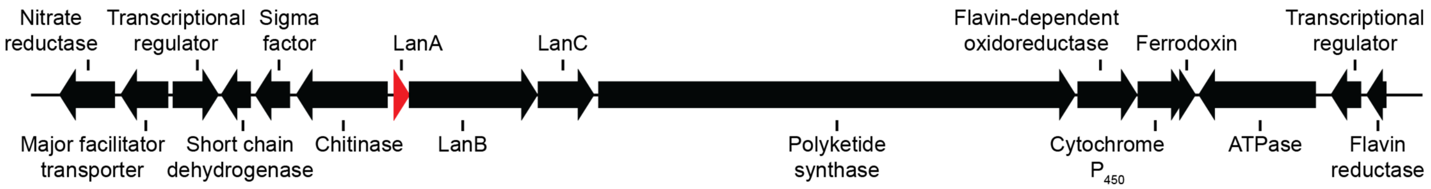


LanA sequence MNNISAAETEFDLDLDLDTAFEDAPAMEAKSFTGTGTCLSCPVTSWHC

#### Class II

*Actinoplanes brasiliensis* strain DSM 43805

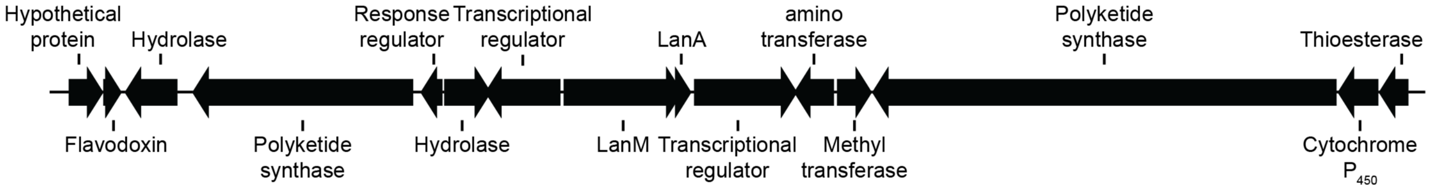


LanA sequence MSEIKNDQNDECGESEQITEHAVGRLRLLPSVSFGTRAAALAQIALPVAVVTAGITAAVHGMGAENLVAI ETSCCPPSPR

#### Class III

*Nocardia* sp. NRRL S-836

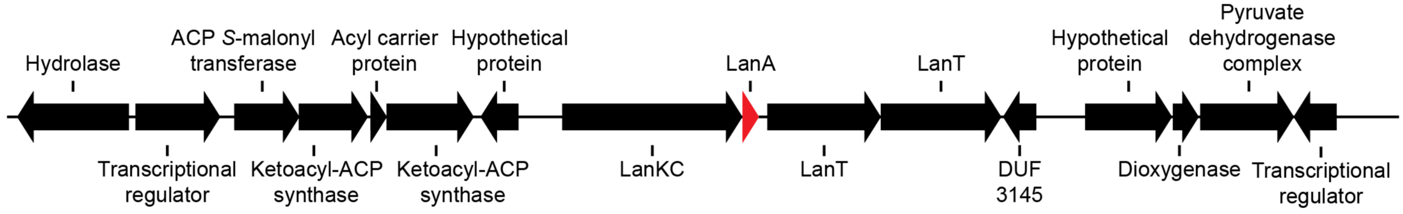


LanA sequence MGFILDMQDLETPEAPANVLAGGSSGGGGGSTTASNASLLLPCSHSTVSLLAC

#### Class IV

*Amycolatopsis jejuensis* strain NRRL B-24427

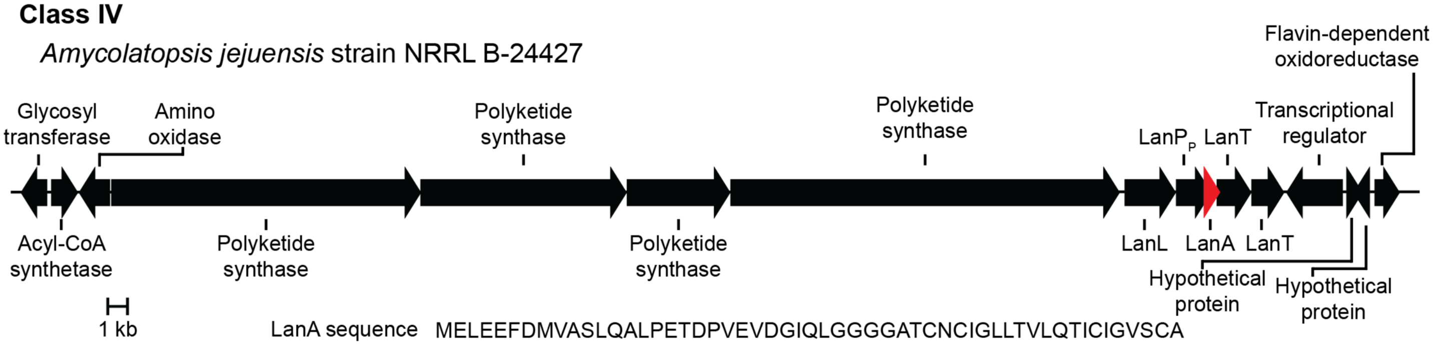


LanA sequence MELEEFDMVASLQALPETDPVEVDGIQLGGGGATCNCIGLLTVLQTICIGVSCA

**Supplementary Figure S9 continued.**

### Zinc-dependent alcohol dehydrogenase containing clusters

#### Class I

*Streptomyces* sp. CB02959

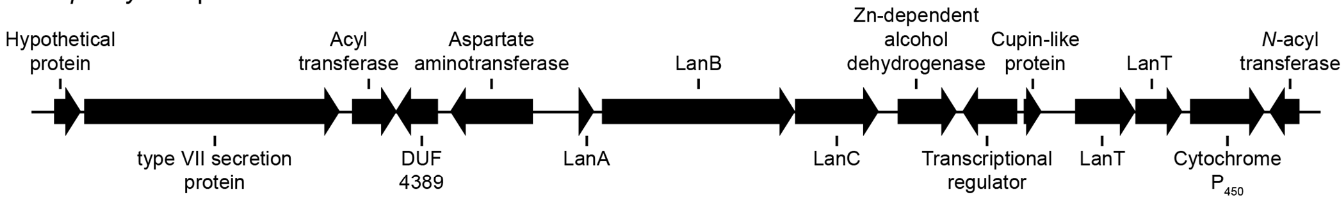


1 kb

LanA sequence MRTEIVLQEFDDADLDLDLRISDVSDQAQEFGQGTYTSPSSYAIGTRCPTCC

#### Class II

*Enterococcus faecalis* strain 4928STDY7071440

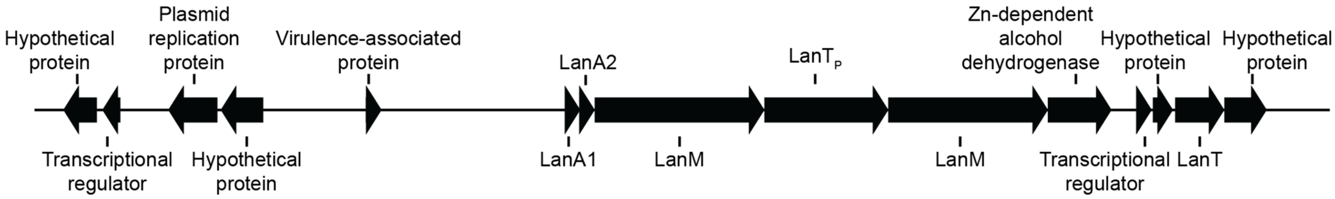


1 kb

LanA1 sequence MRDEENNETLINNEQVAWFEEVADQSFDDDVFGACTTNTFSLSDYWGNKGNWCTATHECMGWCK

LanA2 sequence MNNDKQNEPNVDLELGKYLEADLISLTDDEVTGGGTPTITIPISIAVSGWISDKTCPTSVCTRAC

#### Class III

*Nonomuraea* sp. ATCC 55076

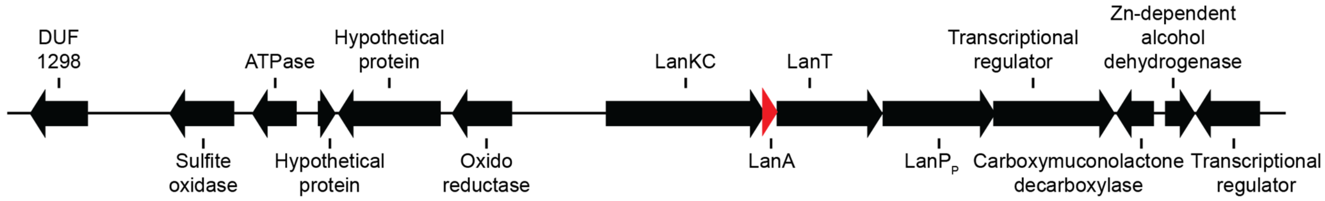


1 kb

LanA sequence MVLLDLQGLEAPAASDVASGGSTLTVLSCHSGKPSNLSVALCH

#### Class IV

*Kitasatospora* sp. CB01950

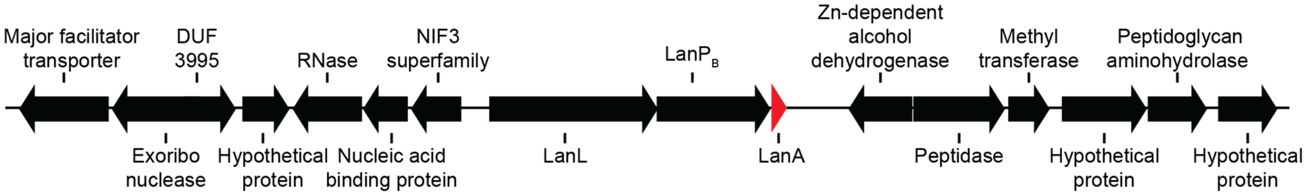


1 kb

LanA sequence MENMTAFETDLAALQELPELESVELGGHGAGCQFTCLVLTCLIFTIS

**Supplementary Figure S10.** Phylogenetic distribution of genomes in the dataset used for this study.

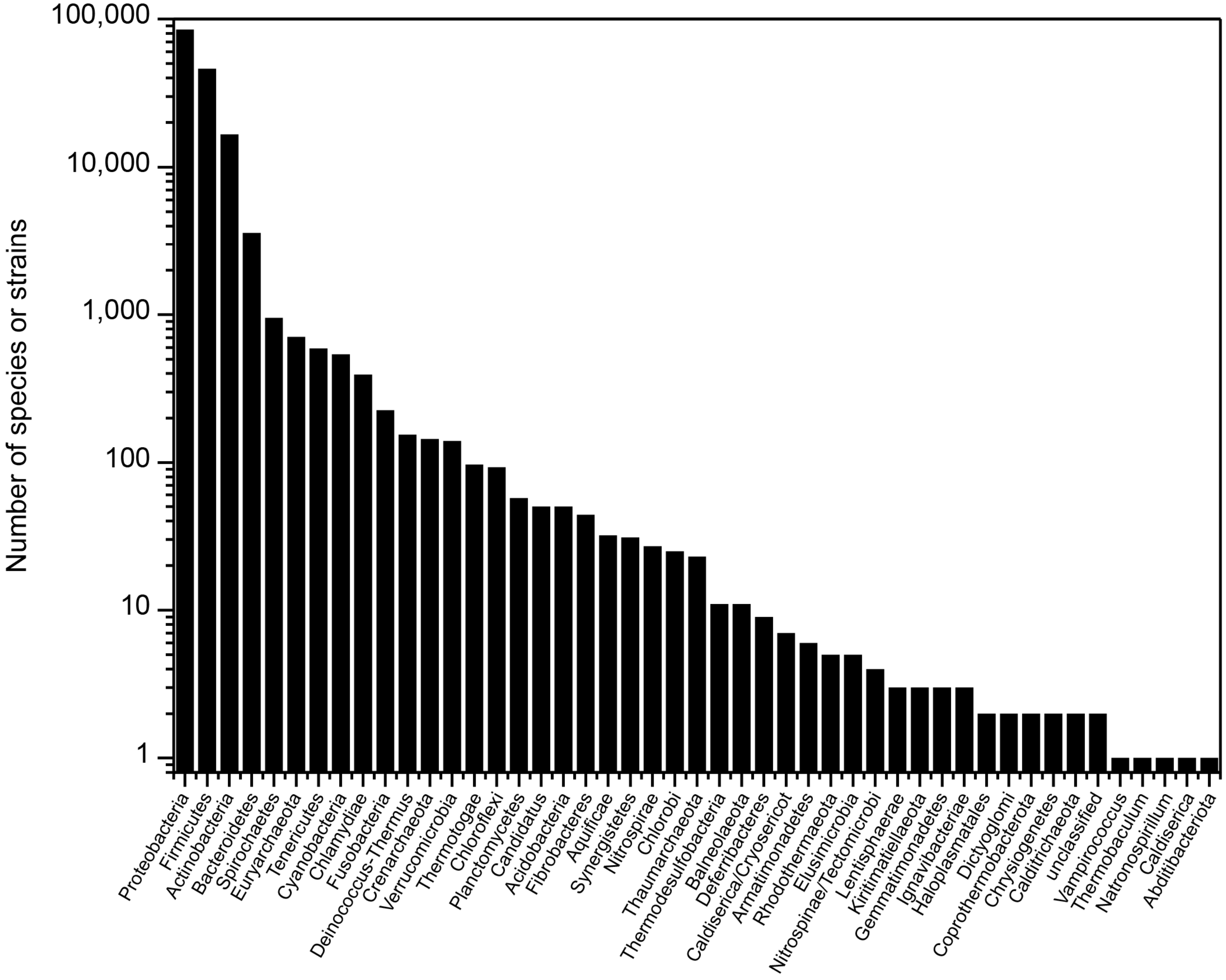

**Supplementary Figure S11.** An approximate maximum likelihood, midpoint rooted phylogenetic tree of LanC and LanC-like domains including human LanC-like proteins. The branches for the human LanC-like proteins are colored in red.

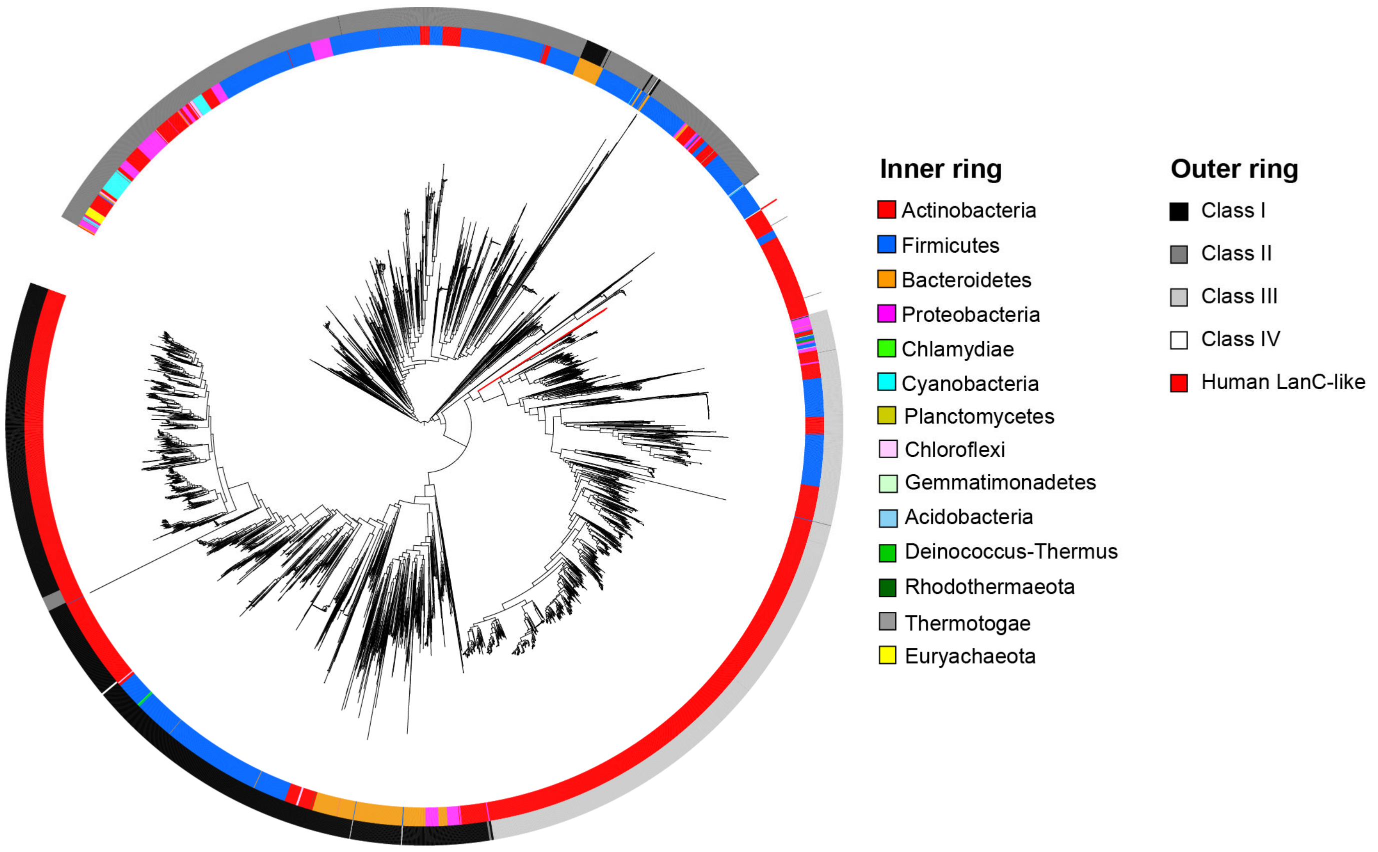

**Supplementary Figure S12.** GC content of clusters versus the cognate genome. The red diagonal line is the linear regression with slope and intercept given in the figure. Blue points are the 10 clusters with the largest difference in GC content between the cluster and the genome. The box plot represents the differences between the cluster GC content and the genome GC content. The boxes are 1 SD and the whiskers are the maximum and minimum values.

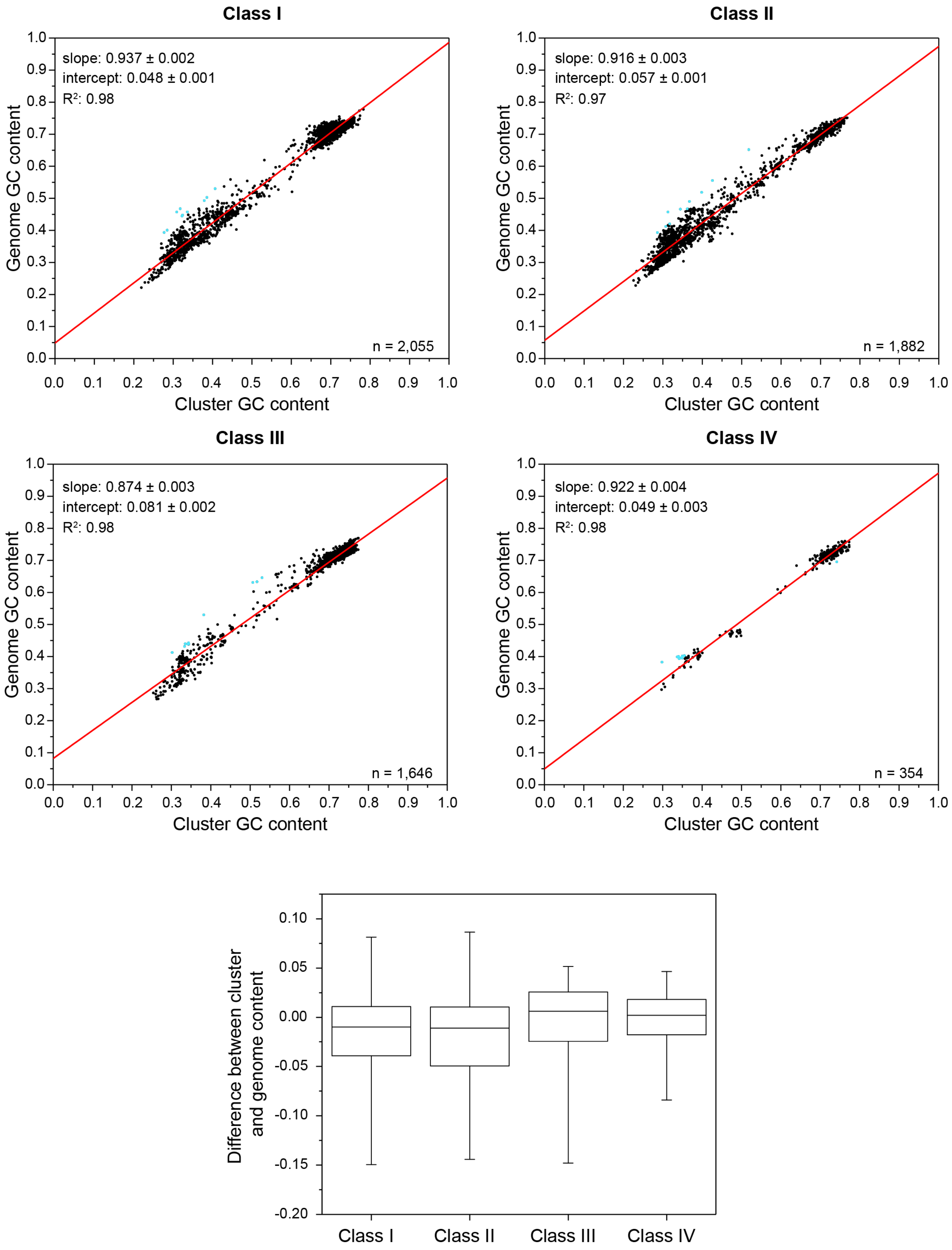
